# Hierarchical Domain Structure Reveals the Divergence of Activity among TADs and Boundaries

**DOI:** 10.1101/361147

**Authors:** Lin An, Tao Yang, Jiahao Yang, Johannes Nuebler, Guanjue Xiang, Ross C. Hardison, Qunhua Li, Yu Zhang

## Abstract

The spatial organization of chromatin in the nucleus has been implicated in many aspects of regulated gene expression. Maps of high frequency interactions between different segments of chromatin have revealed Topologically Associating Domains (TADs), within which most of the regulatory interactions are thought to occur. Recent studies have shown that TADs are not homogeneous structural units, but rather they appear to be organized into a hierarchy. However, precise identification of hierarchical TAD structures remains a challenge. We present OnTAD, an Optimized Nested TAD caller from Hi-C data, to identify hierarchical TADs. Compared to existing methods, OnTAD has significantly improved accuracy and running speed. Results from OnTAD reveal new biological insights on the role of different TAD levels, boundary usage in gene regulation, the loop extrusion model, and compartmental domains. The software and documentation for OnTAD are available at: https://github.com/anlin00007/OnTAD

## Background

Previous studies have shown that the human genome is spatially organized at different levels in the nucleus, with each level of organization playing a role in gene regulation [1]. Starting with the original Chromatin Conformation Capture (3C) assay [2] for measuring chromatin interaction frequencies, many higher throughput, sequencing-based methods such as 4C, 5C, ChIA-PET, Hi-C, and Hi-ChIP have been developed to measure 3D interaction frequencies at different resolutions [3–8]. These maps of interaction frequencies between segments of chromatin are interpreted in terms of chromatin structures. Among the methods, Hi-C [9] obtains measurement of chromatin interaction frequencies across the entire genome. Within some local regions in a genome, interactions are significantly higher than they are to adjacent regions; these highly interacting regions are termed ‘Topologically Associating Domains’ (TADs) [10,11]. The proteins CTCF and cohesin are frequently enriched at TAD boundaries, and they have been implicated in the formation of an isolated local environment [11]. Furthermore, the positions of many TADs are similar across different cell-types and even conserved between species [11,12]. As a result, TADs have been widely interpreted as a basic architectural unit within which many gene regulatory interactions occur. To date, several computational methods have been developed to locate TADs in the genome. For example, Dixon et al. [11] developed a ‘Directionality Index’ based on the shift of interaction direction from upstream to downstream to estimate boundaries of TADs. Other methods, such as TOPDOM [13] and Insulation Score [14], convert the TAD boundary finding problem to a local minimum identification problem by calculating average interaction frequency of surrounding regions at each locus.

While many earlier TAD calling methods treat TADs as a single structure, recent high-resolution studies have shown that TADs contains internal substructures, with sub-TADs nested within larger TADs [15–19]. Several recently developed TAD calling methods aimed to identify nested TAD structures. For example, TADtree [15] identifies TADs based on a relationship between the enrichment of contact frequency and TAD size, and assembles TADs into a TAD tree that best fit the contact matrix. rGMAP [16] assumes that the interaction frequency in sub-TADs is different from those in larger TADs, and applies a Gaussian Mixture model to identify both types of TADs. Arrowhead [17] identifies corners of TADs at multiple sizes, allowing TADs and subTADs to be detected simultaneously. 3D-Net [18] utilizes a maximization of network modularity to identify TADs at different levels. And finally, IC-Finder [19] uses a hierarchical clustering method to identify the TAD hierarchy.

Although the aforementioned methods provide useful tools for identifying TADs and their internal substructure, we still lack comprehensive understanding of the functions of the hierarchical structures within TADs. Recent work on low-resolution Hi-C data [20] has shown that, at the large scale (> 1Mb), TADs can form a hierarchy of domains-within-domains (“metaTAD”) through TAD-TAD interactions, and the successive levels of metaTAD organization correlate with key epigenomic and expression features. This raises the natural question: do the hierarchical levels within TADs also correlate with distinctive functional roles in chromosome organization and gene regulation? However, most existing TAD callers focus on identifying the locations of TADs and subTADs, rather than the hierarchical organization within TADs, making them less suitable for investigating the biological functions of TAD hierarchy. Furthermore, many existing callers are computationally inefficient for high-resolution Hi-C data and often lack a principled approach to choosing algorithmic parameters [16]. These issues limit the utility of existing TAD callers for investigating finer TADs structures using high-resolution data.

We present OnTAD, an Optimized Nested TAD caller that efficiently and robustly uncovers hierarchical TAD structures from Hi-C data. Our approach first identifies candidate TAD boundaries by scanning through the genome with a sliding window at a series of different window sizes, using an approach inspired by TOPDOM [13]. Then, the candidate boundaries are assembled into the optimized hierarchical TADs structures using a recursive dynamic programming algorithm based on a scoring function. Our systematic evaluation shows that OnTAD substantially outperforms existing TAD callers for both TAD boundary identification and hierarchical TAD assembly. Using OnTAD, we uncovered novel insights on the potential biological functions of TAD structures. In particular, we observed that active epigenetic states are substantially more enriched in inner TADs than in outer TADs. OnTAD results revealed two categories of TADs, those with or without hierarchical structures, that appear functionally distinct. Compared to nonhierarchical TADs, the boundaries of TADs with hierarchical structures show a higher CTCF enrichment, more active epigenetic states, and a higher level of gene expression. In addition, we observed an apparent asymmetry in TAD boundary sharing, supporting the asymmetric loop extrusion model for the formation of TADs [21]. Together, these results demonstrate that OnTAD is a powerful tool for inferring different levels of chromatin organization across a genome in high-resolution Hi-C data, which should facilitate improved investigations into the roles of chromatin organization in gene regulation.

## Results

### The OnTAD algorithm

OnTAD takes a Hi-C contact matrix as the input and calls TADs in two steps. In the first step, the method finds candidate TAD boundaries using an adaptive local minimum search algorithm inspired by TOPDOM [13]. Specifically, it scans along the diagonal of a Hi-C matrix using a *W* by *W* diamond-shaped window (Figure 1a), calculating the average contact frequency within each window. The locations at which the average contact frequency reaches a significant local minimum (1.96 standard deviations less than local maximum) are identified as candidate TAD boundaries (see Methods). Because the sizes of TADs are unknown, OnTAD repeats the above steps using a series of window-sizes, *W*= 1,2.,…,*K*, to uncover all possible boundaries for TADs in different sizes. Here, *K* depends on the resolution of the Hi-C matrix and the maximum TAD size that the user aims to call. For instance, for a 10kb resolution Hi-C matrix and a maximum TAD size of 2Mb, *K*=2000/10=200. The union of the candidate boundaries of all window sizes is used to assemble TADs in the next step (Figure 1b).

**Figure 1.**
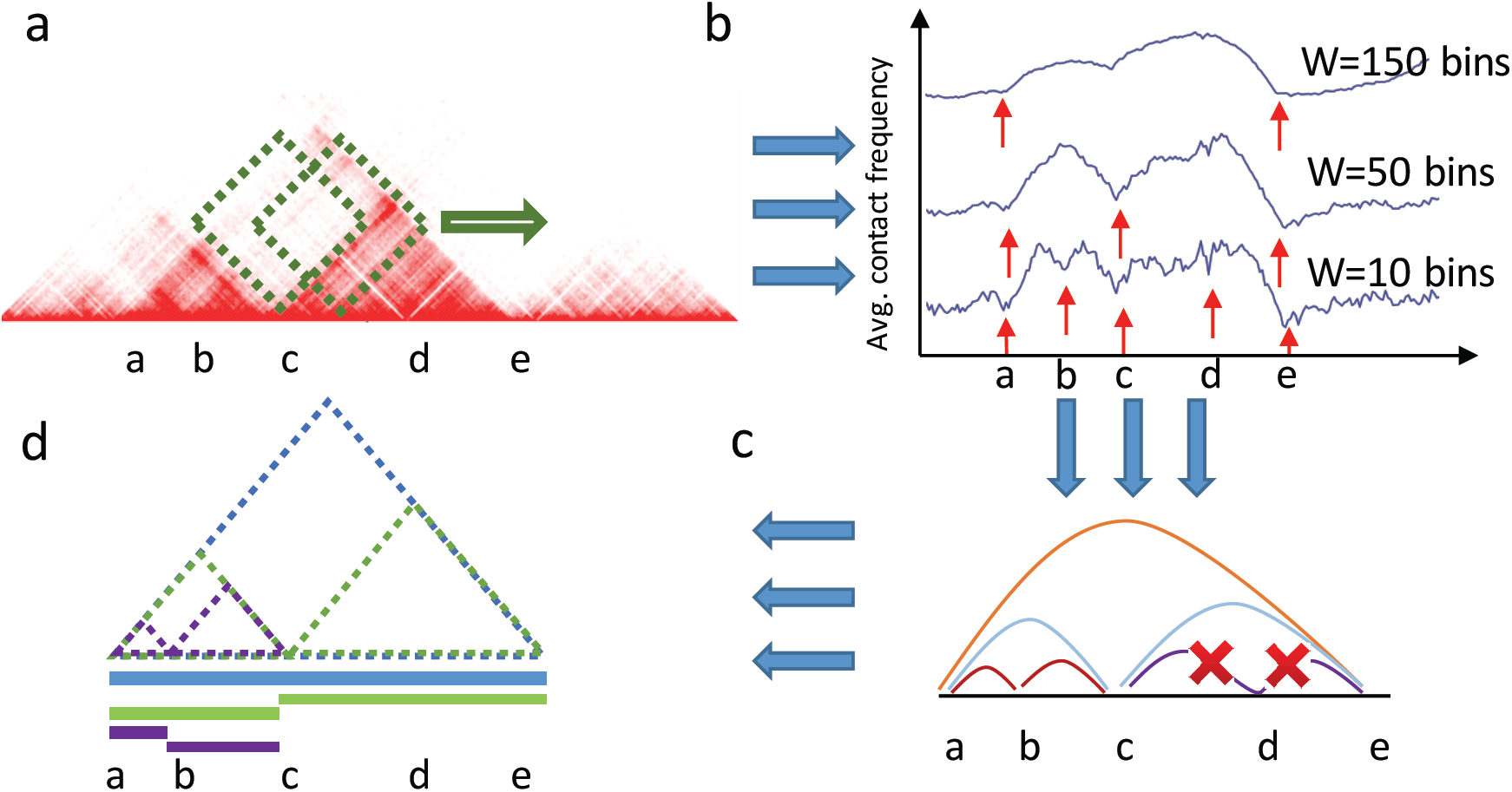
Overview of the OnTAD pipeline. **a**, OnTAD uses a sliding diamond-shaped window to calculate the average contact frequency within the window at each locus on the genome. The five loci marked by letters ‘a’-’e’ are examples being evaluated as potential TAD boundaries, with ‘d’ being a clear false positive. **b**, Identification of candidate TAD boundaries in OnTAD. Blue curve: the average contact frequency of the diamond-shaped windows, calculated at different window sizes (W) and different loci. Red arrows: the location of significant local minimums of the average contact frequency, i.e. candidate TAD boundaries. **c**, OnTAD assembles candidate boundary pairs using a Dynamic Programming algorithm (see methods) **d**, Visualization of the final output from OnTAD. In the genome browser, the identified hierarchical TAD is displayed as a series of horizontal bars, where each (sub)TAD is represented as a horizontal bar colored according to its TAD level.

In the second step, OnTAD assembles TADs by selectively connecting pairs of candidate boundaries using a dynamic programming algorithm (see Methods). To form a TAD between a pair of boundaries, OnTAD requires the mean contact frequency within the potential TAD area between the boundaries to exceed that of the surrounding area outside of the TAD by a user-defined margin (*λ*); otherwise, no TAD is formed between the boundaries. The dynamic programming algorithm is formulated to recursively identify the optimal partition of the genome for yielding the largest rightmost subTADs within each identified TAD according to a score function (Supplementary Figure 1) that de-convolutes the contact frequency signals across the TAD hierarchy (Supplementary Figure 2). At the end of the recursive procedure, the optimized solution that maximizes the score function is obtained (defined in Methods), producing a hierarchical TAD organization that best fits the observed Hi-C contact matrix. The locations of the identified TADs are provided to the users as a plain text file and a bedgraph file ready for visualization on genome browsers.

### Comparison with existing TAD calling methods

We compared OnTAD with four representative TAD calling methods (DomainCaller, rGMAP, Arrowhead and TADtree) using the Hi-C data in GM12878 from Rao et al.[17]. Each method was run using the settings recommended in its manual (see Supplementary file 1 for the version and parameters for each method). All the evaluations were performed using 10kb resolution for the normalized genome-wide Hi-C data, unless specified otherwise. We also tested OnTAD on raw data, and the results obtained were similar to those observed for normalized data (Supplementary Figure 9).

#### Accuracy of TAD boundary detection

We first evaluated the accuracy of TAD boundary detection using enrichment of architectural proteins at boundaries as a reference for accuracy. CTCF is an architectural protein implicated in formation of TAD structures [10]. Thus, we expect a high concentration of CTCF signal (from ChIP-seq data) at accurately called TAD boundaries. We computed the average CTCF ChIP-seq signal in the boundaries identified by each TAD calling method as well as their neighborhood regions. As shown in Figure 2a (left panel), all methods showed enrichment of CTCF signal in the identified TAD boundaries over that in the surrounding regions (fold change > 1.63). Among them, OnTAD had the highest CTCF enrichment (mean signal 1.22X greater than that of the second highest method, T-test p-value = 1.91e-28). A similar result was obtained for the enrichment of the RAD21 and SMC3 subunits of the cohesin complex, which are also key components in the formation of TADs [21]. The boundaries identified by OnTAD showed a higher enrichment than those identified by other methods (mean signal 1.14X and 1.04X greater than that of the second highest method, T-test p-value = 9.34e-16 and 1.91e-12, respectively) (Figure 2a middle and right panel). The stronger enrichment of CTCF and cohesin signals suggests that OnTAD produces more accurate calls of TAD boundaries than the other methods.

**Figure 2.**
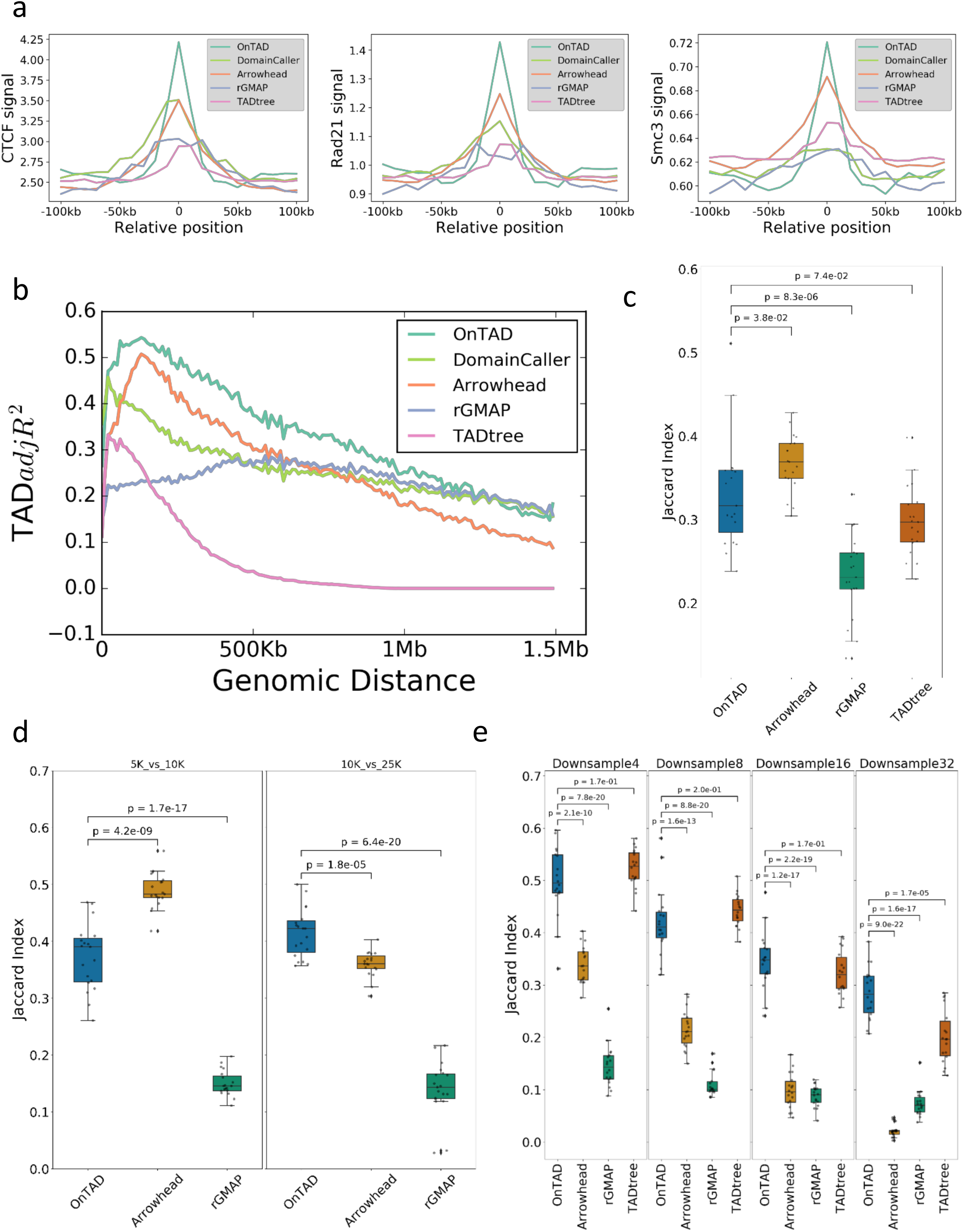
Evaluation of TAD calling methods. **a**, Average ChIP-Seq signal at TAD boundaries and surrounding regions (+/− 10 bins) (from left to right, CTCF, SMC3 and RAD21). **b**, Proportions of Hi-C signal variability explained by the called TADs (measured by TAD-adjR^2^) at different genomic distance between two interacting loci. (Average TAD-adjR^2^: OnTAD: 0.33, Arrowhead: 0.26, DomainCaller: 0.26, rGMAP: 0.23 and TADtree: 0.06). **c-e**, Reproducibility of TAD boundaries (Jaccard index): **c**, between two biological replicates (GM12878, 10Kb) **d**, between resolutions (5Kb vs 10Kb) and (10Kb vs 25Kb). **e**, across different down sampled sequencing depths (GM12878, original vs 1/4, 1/8, 1/16 and 1/32 of the original sequencing depth, raw data was used). Note: TADtree was not included in **d**, because it has difficulty handling data with 5Kb resolution due to its large memory consumption. It also has difficulty for chr1-3 at 10Kb resolution either. Thus these three chromosomes were excluded for all TAD callers in all comparisons.

#### Accuracy of TAD assembly

We next evaluated the accuracy of TAD calling. If TADs are accurately called, one would expect that a high proportion of the variation in the contact frequencies in the Hi-C matrix is explained by TAD calls. We developed a metric called TAD-adjR^2^, which is a modified version of the adjusted R^2^ (see Methods), to measure the proportion of Hi-C signal variation explained by TAD calls. Because contact frequencies decay over the genomic distance between a pair of interacting loci, we stratified the contacts by their genomic distance and calculated TAD-adjR^2^ within each stratum. As shown in Figure 2b, OnTAD has a higher TAD-adjR^2^ than that of the other methods across almost the entire span of genomic distances examined (0-1.5Mb) (Average TAD-adjR^2^: OnTAD: 0.33, Arrowhead: 0.26, DomainCaller: 0.26, rGMAP: 0.23 and TADtree: 0.06). This high level of explained Hi-C variance indicates that OnTAD produces a better classification between TADs and non-TAD regions compared to other methods.

#### Reproducibility of TAD calls and boundaries

Another important criterion for TAD calling is the reproducibility of the identified TADs and their boundaries. To measure the reproducibility of TAD boundaries, we calculated the agreement of boundaries (Figure 2c-e) between two TAD calling results using the Jaccard index. To measure the reproducibility of TADs, we treated each region covered by a TAD as a cluster of bins in the genome, and then measured the agreement of cluster assignments between two TAD calling results using the adjusted rand index (Supplementary Figure 3a-c). We evaluated the reproducibility in three scenarios: 1) between biological replicates (GM12878, 10Kb) (Figure 2c, Supplementary Figure 3a); 2) across different resolutions (5Kb, 10Kb, 25Kb) (Figure 2d, Supplementary Figure 3b); and 3) at different sequencing depths (original sequencing depth versus 1/4, 1/8, 1/16 and 1/32 of the total number of reads) (Figure 2e, Supplementary Figure 3c). As shown in Figure 2c-e and Supplementary Figure 3b, both the boundaries and the TADs identified by OnTAD were fairly reproducible, consistently having either the highest or the second highest Jaccard index or Adjusted Rand index in all scenarios.

#### Run time comparison

We recorded the run time of different methods on the same high-performance computing cluster (Xeon E5-2680CPU and 72Gb RAM). OnTAD ran notably faster than all the other methods (Supplementary Table 1). For example, it took OnTAD 655 seconds to analyze 10Kb resolution data for the whole genome, which was 3X faster than Arrowhead, 24X faster than DomainCaller, 28X faster than rGMAP, and 263X faster than TADtree.

### Level of TAD hierarchy is related to gene activity and epigenomic states

We systematically studied the biological features of the TAD hierarchy, again using the Hi-C data in GM12878 from Rao et al.[17]. Overall, 75.7% of the genome was covered by the TADs identified by OnTAD; the rest of the genome was not assigned to any TADs, and we refer to these TAD-free regions as gaps. Among all TADs identified by OnTAD, the majority (92.2%) contained or belonged to hierarchical structures, while a small fraction had no hierarchical structure. We referred to the former as ‘hierarchical TADs’ or ‘nested TADs’, and the latter as ‘singletons’ (Figure 3a). We hypothesized that chromatin organized into these two types of TADs may be playing distinctive roles in regulation, and thus we examined their association with various epigenetic marks.

**Figure 3.**
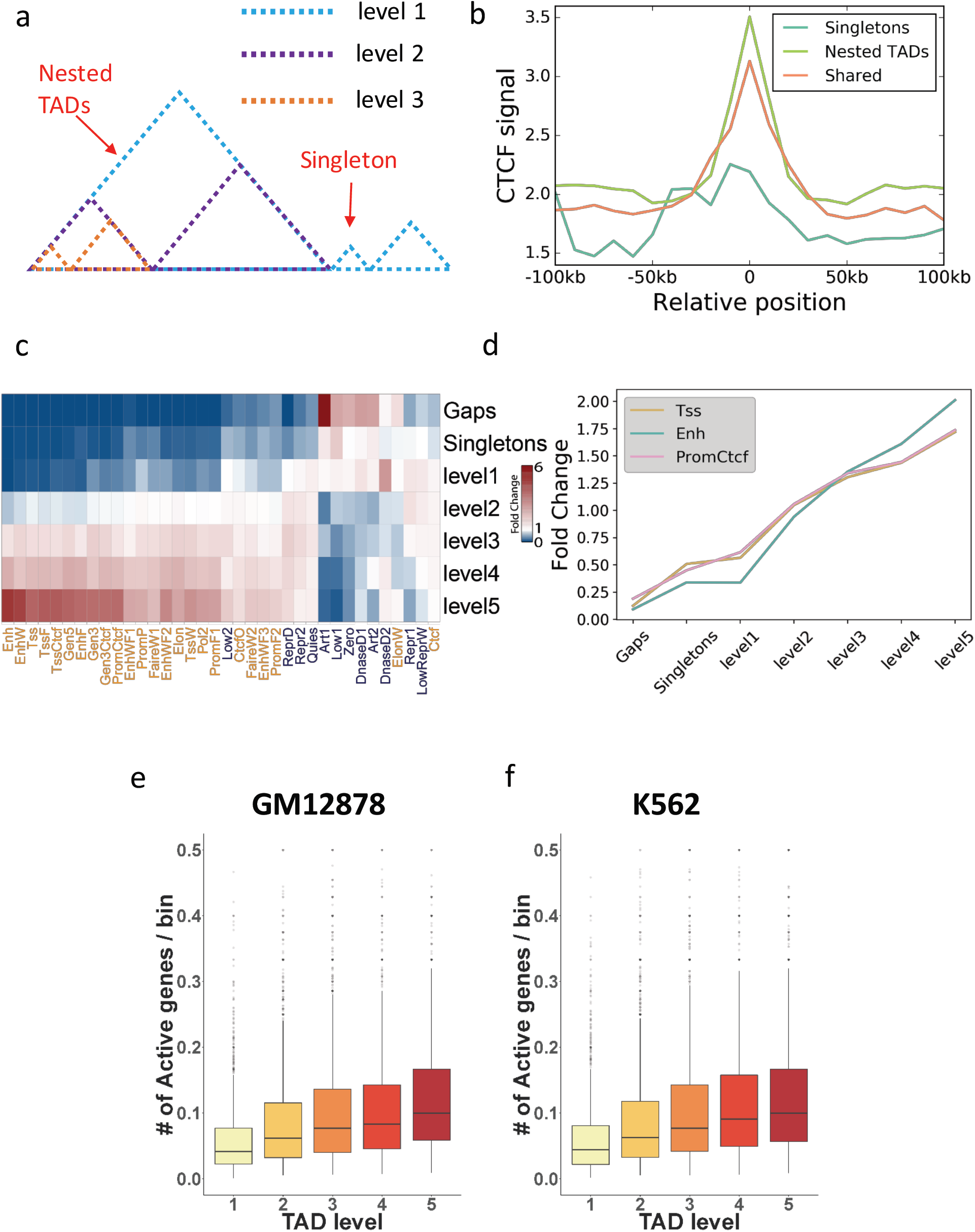
Hierarchical TADs are more active than singletons. **a**, An illustration of hierarchical levels of TADs. The levels are assigned from external to internal. The TADs covered by cyan dash line are assigned to level 1, by blue dash line are assigned to level 2, by orange dash line are assigned to level 3, and singletons are also assigned to level 1 (cyan). **b**, mean CTCF signal at the boundaries specific to hierarchical TADs (light green), specific to singletons (cyan), and shared between hierarchical TADs and singletons (orange). The boundaries of hierarchical TADs have the highest enrichment of CTCF signal. **c-d**, Enrichment of epigenetic states at the regions covered by different levels of TADs. The enrichment (fold change) of active states (marked in orange in **c**) increases as the TAD level increases. The trend is visualized for the states of Tss, Enh and PromCtcf in **d.** The whole-genome average is used as the background for calculating enrichments. **e-f**, Density of expressed gene in different levels of TADs in GM12878 (**e**) and K562 (**f**).

#### Boundaries of hierarchical TADs have a higher CTCF enrichment

We first compared the CTCF enrichment (see Methods) at the boundaries of the two types of TADs. Indeed, the boundaries of hierarchical TADs were substantially more enriched with CTCF signal than singleton boundaries (Figure 3b) (mean CTCF signals are 3.51 and 2.25, respectively, T-test p-value = 1.52e-18). This enrichment of CTCF signal arose from a higher average number of CTCF peaks at the boundaries of hierarchical TADs. The mean number of CTCF peaks per boundary of a hierarchical TAD was 0.451, whereas it was only 0.181 for boundaries of singleton TADs (T-test p-value = 4.52e-30).

#### Hierarchical TADs have a stronger association with active epigenetic states

Chromatin interactions are strongly associated with local, active epigenetic profiles [12,17,22]. We thus expected to observe a positive association between the enrichment of active epigenetic states and the levels of TADs. Starting with the 36 epigenetic states defined by IDEAS segmentation [23] on 6 ENCODE cell lines, we evaluated the association between active epigenetic states and TAD hierarchies. We classified hierarchical TADs into five levels, with level one being the outermost TADs, level two being the immediate subTADs nested under one layer of level one TAD, and so forth until level five, which contains the subTADs nested under four or more layers of TADs in the hierarchy. We observed that the proportion of active epigenetic states increase along the levels of TADs (Figure 3c, d). In contrast, singletons are notably less active compared with hierarchical TADs (especially when level >2). In fact, singleton TADs showed enrichments similar to those for the gap regions. A similar pattern of enrichment for active states in hierarchical TADs was also observed in other cell types (K562 and HUVEC) (Supplementary Figure 4). Taken together, our results showed that hierarchical TADs are on average more active than singletons; and within hierarchical TADs, inner TADs (e.g., subTADs) are more active than outer TADs.

#### Hierarchical TADs have more active gene expression

We further investigated how gene expression is associated with TAD hierarchies. Using the RNA-seq data of GM12878 from the ENCODE consortium (www.encodeproject.org) [24], we defined expressed genes as those with FPKM > 5. Then within TADs at each level, we computed the density of expressed genes (the number of expressed genes per bin, i.e., 10Kb region). If a gene was covered by more than one TAD, we associated it with the innermost TADs. We found that, as the TAD level increases, the density of expressed genes also increases, i.e., genes are more frequently activated within inner TADs than outer TADs (ANOVA test p-value of < 2.2e-16) (Figure 3e). Similarly, we observed the same trend of positive association between density of expressed genes and the TAD level (ANOVA test p-value < 2.2e-16) in the K562 cell line (Figure 3f).

### Shared TAD boundaries are asymmetric and more active than other boundaries

It has been reported that TAD boundaries are interaction hotspots [25]. We also observed that, for TADs at all levels, the number of expressed genes and the enrichment of active epigenetic states are significantly higher at the TAD boundaries than at the internal regions of TADs (all T-test p-values < 0.001) (Supplementary Figure 5a&b). Thus we undertook an additional analysis of TAD boundaries.

We observed that the boundaries of hierarchical TADs were frequently shared by multiple TADs. We hypothesized that the boundary usage may play an important role in maintaining hierarchical structures and regulating gene activities. To investigate this hypothesis, we classified boundaries into five categories, according to the maximum number of TADs that use a boundary on one of the two sides of the boundary (Figure 4a). A boundary is classified as level one if it is used by no more than one TAD on either side, level two if it is used by exactly two TADs on one side and less or equal to two TADs on the other side, and so forth to level five if it is used by five or more TADs on either side. For example, a boundary shared by two TADs to its left and three TADs to its right was classified as level three. The number of boundaries assigned to each category is shown in Supplementary Figure 7.

**Figure 4.**
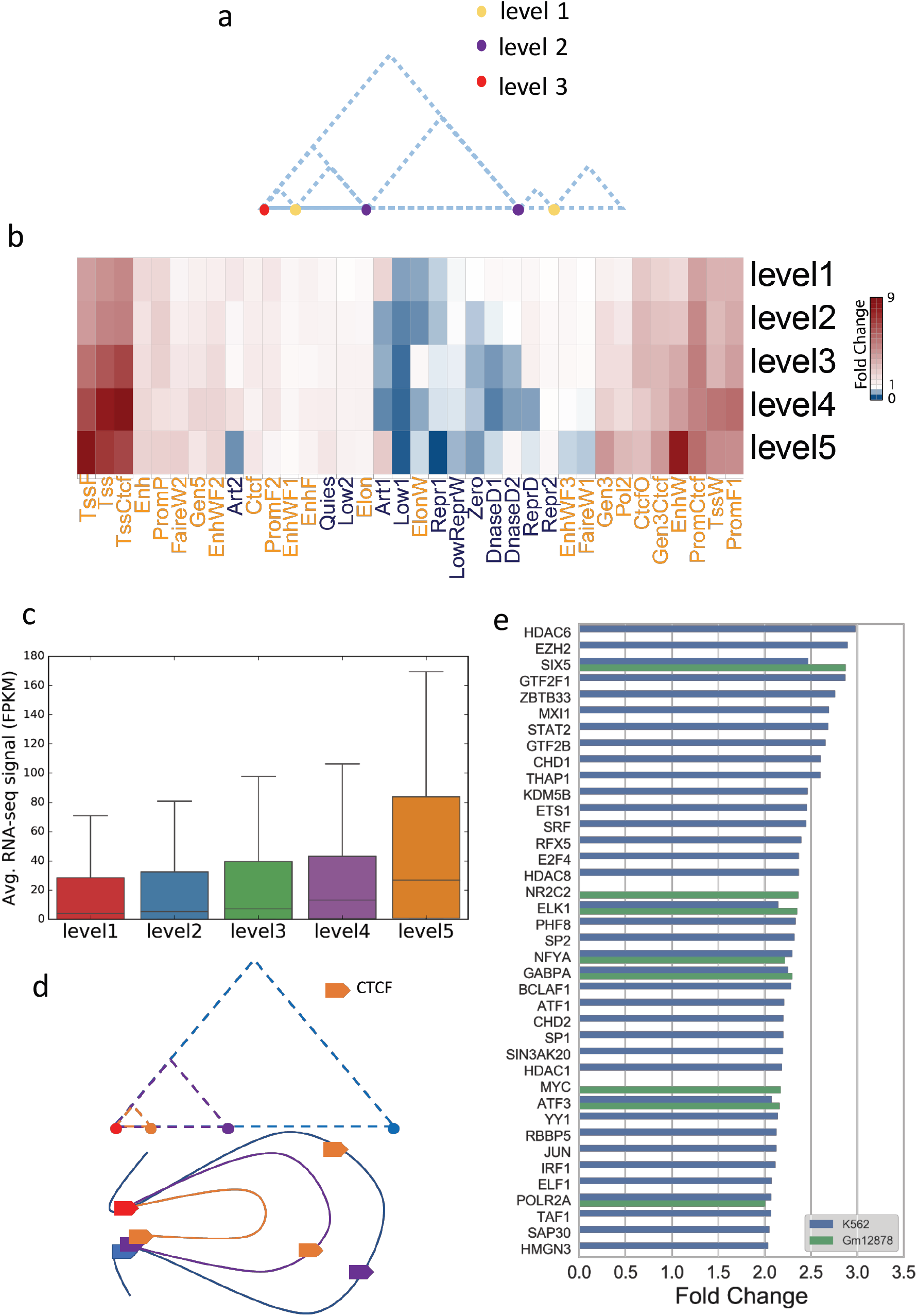
Hub-boundaries are highly active in gene regulation. **a**, An illustration of the TAD boundary levels. The boundary levels are defined as the maximum number of TADs that use a boundary on either its left or right side. The yellow, purple and red dots refer to boundaries of level 1, 2, and 3, respectively. **b**, Enrichment of epigenetic states at different levels of TAD boundaries. Hub-boundaries (i.e. boundaries with level =5) are significantly enriched with Tss related states than others. (Active epigenetic states are marked in orange) **c**, Distribution of gene expression levels for genes whose transcription start sites overlap with TAD boundaries. Genes are classified by the level of TAD boundaries. **d**, Illustration of hierarchical TAD and asymmetric loop extrusion. The red boundary denotes the ‘anchor’ site that starts the loop extrusion in asymmetric loop extrusion model. Boundaries in other colors are the stopping sites of the loop extrusion. The hierarchical TADs are formed by multiple stops of the loop extrusion that share the same start site. **e**, TFs enriched (Fold Change >2) at hub-boundaries in GM12878 and K562 cell lines. The fold change of ChIP-seq TF peaks at hub-boundaries (level =5) against level 1 boundaries is shown.

#### Epigenetic and genomic profiles

We examined the enrichment of active epigenetic states at different boundary levels. We observed a significant positive correlation between the enrichment (fold change) of active epigenetic states and the number of times each boundary is shared (e.g., Tss state: Pearson coefficient = 0.89; TssCtcf state: Pearson coefficient = 0.92) (Figure 4b). We further studied the relationship between gene expression level and boundary sharing. Again, we observed a significant positive association between the number of times a boundary was shared and the gene expression level (ANOVA test p-value = 1.97e-05). In particular, the gene expression level at the boundaries that were shared by 5 or more TADs was substantially higher than that at boundaries that were shared by fewer TADs (Figure 4c). At a higher level boundary, multiple genomic loci (the boundary plus the other ends of the TADs) must be in proximity in three-dimensional space. This situation is reminiscent of chromatin hubs, and thus, we call the boundaries shared by 5 or more TADs “hub-boundaries”. We posited that hub-boundaries are more active in gene regulation than boundaries that are shared by fewer TADs.

#### The asymmetric loop extrusion model

Interestingly, we also observed some asymmetry in boundary usage and TAD formation in hierarchical TADs. Specifically, we have observed (1) a significant difference in boundary usage between the left and the right boundaries of the same outer TAD (Z-test p-value < 2.2e-16) and (2) a significant difference in the numbers of TADs formed by a boundary on its left and right sides (Z-test p-value < 2.2e-16) (Supplementary Table 2).

We therefore asked if the observed asymmetry is related to the mechanism of loop formation. A recent study in yeast suggested that loops are formed in an asymmetric process, where the loop extrusion complex anchors on one side and DNA reels through from the other side [26]. We here hypothesize that loop extruders are preferentially loaded at or near a specific TAD boundary. Then the asymmetric loop extrusion would start from this site on one end, and it could stop at different sites on the other end. Thus, the TADs formed by multiple stops of loop extrusion in this process would all share the anchor site as the boundary on one side, but each has a different boundary on the other side, leading to the observed asymmetric boundary usage (Figure 4d). Another recent study in *Drosophila* Schneider 2 (S2) cells showed that promoters prefer to interact with enhancers downstream of the transcriptional unit [27], leading to a directional preference in TAD formation. Indeed, as shown in Figure 4b, the boundaries shared by multiple TADs are highly enriched with promoters, thus the observed orientation asymmetry in TAD formation around these boundaries could reflect this interaction preference in promoters.

While some proteins (e.g. Ycg1 HEAT-repeat and Brn1 kleisin subunits) have been found to be related to the anchor sites in yeast [28], little is known about the proteins supporting the anchor sites in human. We therefore performed a transcription factor (TF) enrichment analysis using 161 TF ChIP-seq data from the ENCODE consortium [29–31]. By comparing the fold enrichment of each TF signal in hub-boundary (level 5) with the ones at low level (level 1), we found a group of TFs that were highly enriched in hub-boundaries (Fold Change > 2 in either GM12878 (n = 8) or K562 (n = 37)) (Figure 4e). These hub-boundary-enriched TFs were strongly associated with chromosome organization function in Gene Ontology Analysis (FDR = 1.33e-06). They were also shown to be highly connected (p-value < 1.0e-16) in the protein-protein interaction database, STRING (Supplementary Figure 7), suggesting that they may potentially form a protein complex. Together, these results suggest that the enriched TFs may play an important role in forming the anchor sites in the asymmetric extrusion process.

### Hierarchical TAD calling unveils distinct epigenetic features of inner TADs

It has been reported that genomic loci within the same TADs tend to possess similar epigenetic features [22], while loci in different adjacent TADs may show different epigenetic features [17]. However, it remains unclear if the divergence of epigenetic profiles also takes place at the subTAD level. To explore the possible answer to this question, we performed OnTAD on Hi-C data from the mouse G1E-ER4 cell [32]. We observed that the majority of TADs (87.1%) are in active compartments. As shown in the OnTAD genome browser track in Figure 5, the region of (chr19:11.3Mb – 12.2Mb) contains multiple nested TADs. Among them, two adjacent subTADs (11.8Mb – 11.9Mb and 11.9Mb – 12.0Mb) that belong to the same outer TAD (chr19: 11.5Mb – 12.0Mb) exhibit distinct epigenetic features, with enriched repressive epigenetic signal (H3K27me3) in the left subTAD and enriched active epigenetic signal (H3K27ac, H3K4me3 and H3K36me3) in the right subTAD. This demonstrates that, although TADs were traditionally considered to be a fundamental unit of chromatin organization, epigenetic features can be distinctively different between subTADs. Our results show that the subTADs identified by OnTAD better represent homogeneous units associated with epigenetic functions, capturing distinct functional features within subTADs. By identifying these subTADs, OnTAD enables a finer investigation of the hierarchy of chromatin organization and its functionally homogeneous structures.

**Figure 5.**
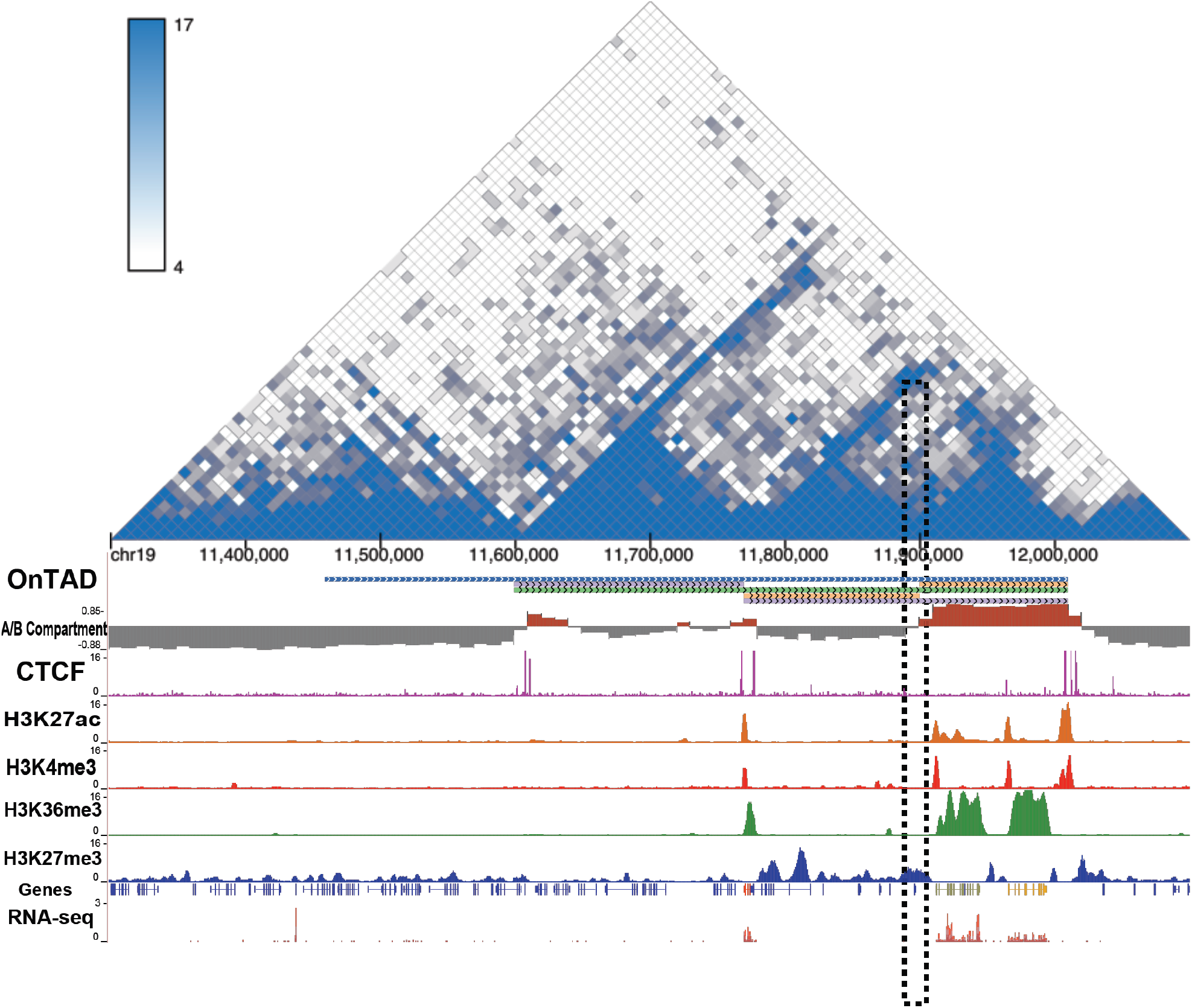
subTADs exhibit distinctive epigenetic profiles. The captured region is chr19:11.3Mb – 12.2Mb in mouse G1E-ER4. The Hi-C heatmap shows a nested TAD structure in this region. OnTAD results are displayed in the genome browser track: blue line denotes level 1 TAD, green line denotes level 2 TAD, purple denotes level 3 TAD and orange denotes level 4 TAD. The two subTADs (orange lines) exhibit distinctive epigenetic features, with one enriched with repressive signals (H3K27me3) and silenced expression (low RNA-Seq signal) and the other enriched with active signals (H3K27ac, H3K4me3, and H3K36me3) and expression (high RNA-Seq signal). The shared boundary (marked by dash box) between these two subTADs has no CTCF peak, indicating the formation of these two subTADs may not involve loop extrusion.

Interestingly, we observed that whereas the outer TAD (11.8Mb – 12.0Mb) had clear CTCF signals at its boundaries, the shared boundary between the two subTADs (11.8Mb – 11.9Mb and 11.9Mb – 12.0Mb) had no CTCF signals from the CTCF ChIP-seq data. This indicates that the formation of these two subTADs is probably independent of CTCF. Furthermore, we performed a high-resolution compartment analysis (See Methods). It showed that the two subTADs fall in different compartments. These results can be interpreted within the framework of recently proposed ‘compartmental domains’ [33,34], which are hypothesized to be formed by A/B compartment without the involvement of CTCF. We will discuss this mechanism further in the discussion.

## Discussion

While hierarchical structures in TAD formation have been reported [15,16,18,19], the involvement of these hierarchies in gene regulation mechanisms remains poorly understood. This is partly due to the lack of a method that systematically identifies TAD hierarchies from Hi-C data and investigates the association of TAD hierarchies with epigenetic features. Here we introduce OnTAD, a new method to uncover the hierarchical TAD structures from Hi-C data. Based on a dynamic programming procedure that recursively finds the best domain partition of Hi-C contact matrix in a hierarchical manner, OnTAD identifies the hierarchy of TADs and their boundaries. It produces a convenient output for visualizing the hierarchy in a genome browser, greatly facilitating the investigation of the interplay between hierarchical TADs and other epigenetic features in gene regulation. Our comprehensive evaluation shows that OnTAD substantially outperforms the existing TAD calling methods in both accuracy and computational efficiency. These results demonstrated the effectiveness of OnTAD for identifying TAD hierarchies and investigating their biological functions.

Using the results from OnTAD, we investigated how hierarchies within TADs were associated with features related to function. In particular, we observed that, on average, hierarchical TADs were significantly more active than TADs without hierarchies (i.e. singletons). The active epigenetic states and active genes were also significantly more enriched in the boundaries shared by multiple TADs (e.g. hub-boundaries) than those used exclusively by a single TAD. These observations echo those on the hierarchy of metaTADs, which also showed a positive association between the enrichments in promotor activity and gene density and boundary usage [20]. Interestingly, we also observed a significant asymmetry in boundary usage between the left and right boundaries in the hierarchical TADs and an asymmetry in the orientation of TAD formation around hub-boundaries, supporting the asymmetric loop extrusion model [26] and preferential orientation of promoter interaction [27].

Our results pose several interesting questions about the mechanisms utilized to form these structures. First, how are these hierarchical TADs structures formed? Are they produced by hierarchical chromatin folding in single cells, or does the nesting reflect a collection of different interaction patterns in individual cells that looks like a hierarchy when the data from a population of cells is aggregated in bulk cell Hi-C data? Single-allele chromatin interactions do reveal regulatory hubs [35], supporting the interpretation that these complex interactions occur in individual cells. A recent single-cell analysis of high-throughput Oligopaint labeling and imaging on Chr21 of A549 cell, showed that both TAD and sub-TAD structures exist in single cells [36]. Furthermore, nested TAD structures could be formed by multi-site interactions in a single cell [36]. However, as acknowledged by the authors, it is still possible that some other domain structures resulted from population averaging. In principle, OnTAD can also be applied to single-cell Hi-C data to explore this question. However, the genome coverage in current single-cell Hi-C data is still low and can only support the analysis at the resolution of ∼100Kb, limiting the detection of finer domain structures (typically ∼50Kb for subTADs we identified). Future studies with higher resolution single-cell Hi-C data will be valuable for addressing this question at a genome-wide scale.

Second, what are the mechanisms to form the hierarchical structures? As observed in our analyses (Figure 5), though the majority of the outer TAD boundaries were bound by CTCF, some subTADs appear to be formed without CTCF binding at their boundaries. The formation of the latter can be explained by the recently proposed ‘compartmental domains’ mechanism [34], which forms domains by establishing A/B compartments without the involvement of CTCF or loop extrusion. Because OnTAD does not rely on CTCF information for TAD identification, it can capture all domain structures, regardless the formation mechanisms. The example in Figure 5 could be explained by joint processes of loop extrusion (for the outer TAD) and establishment of ‘compartmental domains’ [34] for the inner TADs.

In summary, we have demonstrated that the hierarchies of TAD structures are associated with gene regulation and have provided a powerful tool for exploring this association. Though previous results based on low-resolution data suggest that the majority of TAD structures are conservative across cell lines [11], recent analyses found that certain locally frequent interaction regions within TADs are cell type specific [25]. It will be particularly interesting to use OnTAD to systematically investigate how the finer domain structures within TADs differ across cell types, for example, how the levels of hierarchy differ across cell types, and how the changes in hierarchy are associated with differential gene regulation. The biological insights generated by analyses of the finer domain structures should help improve our understanding of the role of chromatin conformation in gene regulation.

## Methods

### Notations and data preprocessing

Let *X* denote a symmetric Hi-C matrix, where each entry (*i,j*) in the matrix is a value quantifying the strength of the chromatin interaction of between bins *i* and *j*. The Hi-C matrix can be raw contact matrix or the normalized matrix produced by the normalization procedures such as ICE [37] and KR [17]. Let *X*[*a:b, c:d*] = {(*i,j*): *a* ≤ *i* ≤ b, *c* ≤ *j* ≤ d} denote a sub-matrix of *X*. A candidate TAD between bins *a* and *b* corresponds to a diagonal block matrix *X*_*[a,b]*_=*X*[*a:b, a:b*], where the mean of the entries in *X*_*[a,b]*_ is expected to be higher than that in its neighboring matrices. Because of the distance dependency in Hi-C data, i.e., the dependence of contact frequency on the proximity of the interaction loci, we normalize the Hi-C matrix before TAD calling by subtracting the mean counts at each distance.

### Identification of candidate TAD boundaries

We identify candidate TAD boundaries using a procedure motivated from the TOPDOM method [13]. This procedure scans the diagonal of the Hi-C matrix, using a sliding square submatrix whose bottom corner locates on the diagonal (Figure 1a), and computes the mean Hi-C signals covered by the submatrix at each location, which is the TOPDOM statistic in [13]. As shown in [13], when the corner of the submatrix lands on a TAD boundary, the TOPDOM statistic reaches a local minimum. Thus, the local minimums of the TOPDOM statistic can be used as candidate boundaries. The original TOPDOM paper only computed the statistics at a fixed window size. To identify all candidate TAD boundaries for TADs in different sizes, the TOPDOM statistics are calculated at all window sizes (W), ranging from 1 to a maximum TAD size (*d*) specified by users. Here, we set the minimum size =3 bins, because structures smaller than 3 bins are too small to form a domain. We set the maximum size=200 for 10kb Hi-C data, because TADs are known to be smaller than a few Mbs.

For each window size W, we first obtained a set of local minimums of the TOPDOM statistics, which are defined as the smallest value in the neighborhood of [i-Lsize, i+Lsize]. To reduce false positives due to noise, the local minimums that are not significantly smaller than the local maximums in the same neighborhood are pruned. Here we required the local minimums to be at least 1.96S smaller than the local maximums to be qualified as a candidate boundary, where S is the standard deviation of the TOPDOM statistic in the entire matrix. The parameter 1.96 is chosen based on the 95% confidence level of a normal distribution, which reasonably approximates the distribution of TOPDOM scores.

Figure 1b shows examples of the local minimums on the genome at different window sizes. Because different window sizes capture the information of TADs in different sizes, we took the union of the pruned local minimums over all window sizes, and used the corresponding bins as candidate TAD boundaries. We selected z according to the procedure described in the section of Parameter Selection. For all the analyses in this work, we used Lsize = 5. It can be adjusted by users.

### Recursive TAD calling algorithm

We developed a TAD calling algorithm to assemble TADs from the candidate boundaries. Several issues need to be considered in the design of the algorithm in order to produce biologically meaningful TADs. First, because a region may be shared by multiple TADs, the scores of these TADs can be strongly correlated. Second, in the TADs with nested structures, the scores of the TADs and their nested sub-TADs are convoluted. Third, some boundaries may be shared between TADs. Last, the algorithm needs to be computationally efficient to call TADs in the genome scale.

To address these issues, we developed a recursive algorithm to identify the TADs that give the optimal partition of the genome according to a scoring function g(X) related to the strength of Hi-C signals (see the next section). Our algorithm assumes that any given two TADs are either disjoint (but can share one boundary) or nested (i.e. one TAD is completely within the other). This assumption is required for the dynamic programming to find an optimal solution in polynomial time. While this assumption sometimes may not be true, it greatly reduces the complexity of the problem while still enabling us to 1) de-convolute nested TAD structures, 2) impose shared boundaries, and 3) obtain an efficient algorithmic solution. Our evaluation showed that the majority of the genome follows this assumption (see the subsection below). Even when it is violated, i.e., the boundaries of the TADs cross each other, our method can still produce a reasonable approximation (Supplementary Fig.1C).

Briefly, the algorithm works as follows. Given a matrix *X*_[*a,b*]_, the algorithm starts at the root level to first find the best bin *i* (*a*≤*i*<*b*) to partition the matrix into two submatrices, *X*_[*a,i*]_ and *X*_[*i,b*]_, such that *X*_[*i,b*]_ is the largest right-most TAD in *X*_[*a,b*]_. Since *X*_[*a,i*]_ and *X*_[*i,b*]_ are disjointed, the TADs within each submatrix can be called separately in a recursive manner. At each recursive step, the parent matrix is partitioned into two sub-matrices, and TADs are called within each sub-matrix using the same recursive formula (Supplementary Fig.2A). The recursion stops when *i*=*a*, i.e., the sub-matrix *X*_[*a,i*]_ contains no TAD. After a recursive step is completed, it identifies the best TADs in the current branch according to the scoring function, de-convolutes the TAD signals in the parent matrix by removing signals of inner TADs, and evaluates if the parent matrix itself is a TAD. This process is repeated until the recursion returns to the root level (Supplementary Fig.2B). Note that, because every TAD is the largest right-most TAD of a parent matrix in a recursive branch, this recursive procedure guarantees to traverse all TADs, even though only the largest right-most TAD is called at each step.

#### Evaluation of the violation of the hierarchical TAD assumption

To investigate the frequency of the violation of the hierarchical TAD assumption, we ran OnTAD on high resolution (10Kb) in-situ Hi-C data in GM12878. We segregated regions around the corner of each TAD into four 5*5 quadrants and calculated the average contact frequency of each quadrant (Supplementary figure 8). If this assumption holds, the interaction frequency is expected to be high in the quadrant within TAD (quadrant 1) and relatively low in at least one of the two quadrants (2 & 3) on the two sides outside of TAD corner. As shown in the Supplementary figure 8, the mean frequency patterns of the four quadrants for most of the TAD corners are consistent with our expectations. This suggests that this assumption holds for a majority of the genome. The violation can be remedied by removing the signals from the called TADs and then rerunning OnTADs on the de-clumped Hi-C data to identify additional TADs.

### The scoring function

Our scoring function *g*(*X*_[*a,b*]_) for matrix *X*_[*a,b*]_ is defined as

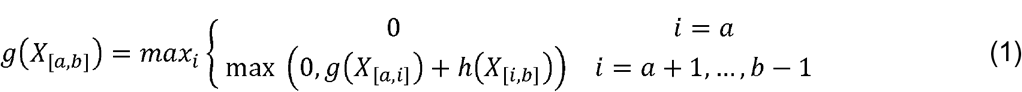

where *h*(*X*_[*i,b*]_)= *g*(*X*_[*i,b*]_)+ Δ(x_[*i,b*]_|sub TADs)

Here, *g*(*X*_*[a,b]*_) is the score of TADs within *X*_[*a,b*]_, not including the score for *X*_[*a,b*]_ itself being a TAD. It is calculated by finding the best left boundary of the largest right-most TAD in *X*_[*a,b*]_. *h*(*X*[*i,b*]) is the score of the largest right-most TAD in *X*_[*a,b*]_. It is the sum of the score of TADs within *X*[*i,b*] and the score of *X*[*i,b*] itself being a TAD, namely Δ(*X*[*i,b*]|sub TADs). For any diagonal block matrix to be called a TAD, its mean signal is required to be greater than the means of its neighboring regions on both sides. We therefore define

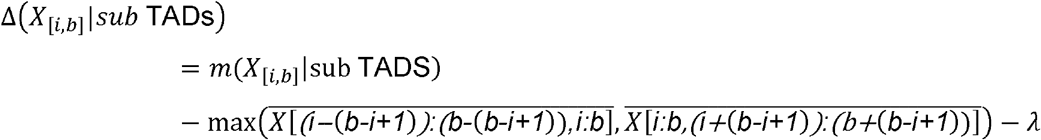

where *m*(*X*_[*i,b*]_|sub TADs) denotes the mean of *X*_[*i,b*]_, excluding the TADs within *X*_[*i,b*]_, returned by the recursion; *λ* is a user-specified nonnegative penalty parameter; *X*[(*i*−(b−i+1)):(b−(b−i+1)),*i*:*b*] and *X*[i:b,(*i*+(*b*−*i*+1))] are two (b−i+1)-by-(b−i+1) off-diagonal matrices in the adjacent flanking regions of *X*_[*i,b*]_; and finally, and 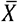 denotes the mean of *X*. We note that Δ(*X*_[*i,b*]_ | *sub* TADs) is calculated based on the TADs returned from *g*(*X*_[*i,b*]_). That is, we do not directly optimize *g*(*X* _[*i,b*]_)+ Δ(*X*_[*i,b*]_| *sub* TADs). The parameter *λ* serves as a threshold for TAD calling. That is, a TAD will be called only when the mean contact frequency within the potential TAD area between the boundaries exceeds that of the surrounding area outside of the TAD by the margin of *λ*. The procedure for selecting *λ* is described in Parameter Selection.

When the score of a candidate TAD is <0, it is likely not a real TAD. We therefore set a lower bound on the score at 0 and do not output the “TAD” with a score 0.

### Parameter Selection

We selected the value of *λ* based on the False Discovery Rate (FDR) of TADs identification. The FDR is calculated as follows. First, the entries in the real Hi-C matrix are permuted within each genomic distance. This results in a null Hi-C matrix that has the same marginal signal distribution as the original Hi-C matrix but without biologically meaningful TAD structures. Next, OnTAD is run on both the original and the permuted Hi-C matrix for a series of *λ*. The TADs identified from the original Hi-C matrix are treated as ‘discoveries’ (R), which is a mixture of false and true discoveries, and those from the permuted Hi-C matrix are treated as ‘false discoveries’ (V), which is used to approximate the proportion of false discoveries in R. Recall that OnTAD assigns each TAD a score according to the scoring function (1). Given a TAD size, the magnitude of the score reflects the strength of evidence to call TAD. Because larger TADs tend to have a lower mean contact frequency after removing their inner TADs, the score is usually smaller for larger TADs. Therefore, we computed the FDR accounting for TAD size. Specifically, for a given value of *λ*, the identified TADs are first stratified by their sizes and scores. Let n be the total number of TADs identified on the original matrices, and *R*_*i*_ and *V*_*i*_ be the numbers of TADs in the th stratum from the original and the permuted matrices, respectively. Then if a TAD (j) is in the th stratum, the probability for the TAD to be a false discovery (a.k.a. local false discovery rate [38] is

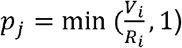

The overall FDR for the TAD identification is computed as the average of the probability to be a false discovery over all TADs identified on the original matrix, based on the relationship between local fdr and FDR [38]:

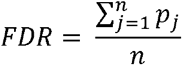

The above FDR calculation is repeated for each value of λ, and the λ corresponding to the FDR cutoff of 0.05 is selected.

In our analysis, the stratum is formed by dividing the TAD calls into 25 equal shares according to the ranking of TAD size (or TAD score, respectively) on the real matrix. This leads to 25*25=625 strata in total. As shown in Supplementary table 3, the FDR is close to 0.05 at λ = 0.1 for GM12878 dataset (10kb). To test the robustness of the tuned parameter, we also performed the same procedure on the mouse G1E-ER4 Hi-C data from Hsu et al. [32] at 10Kb resolution. The FDR was also controlled at the 0.05 level when λ = 0.1 (Supplementary table 4). Therefore, we used λ = 0.1 as the default value in our analyses. In the OnTAD software, we allow users to specify the value of λ to offer more flexibility.

Another important tuning parameter is Lsize, which is the span of the interval (i.e. the interval size = 2*Lsize +1) for searching local minimums of the TOPDOM statistics. This parameter affects the selection of candidate boundaries. If Lsize is too large, some potential boundaries will be missed. If Lsize is too small, the candidate boundary set may include many false positives, increasing the computational burden for the assembly step and the quality of final results. We chose Lsize in the similar way as for choosing *λ* on GM12878 data (10kb). Specifically, we ran OnTAD for different values of Lsize (range=3-10), corresponding to the interval size of 7-21 bins. We chose this range because it is sufficient to cover various TAD sizes. As shown in Supplementary table 5, Lsize=5 (i.e. interval size=11) renders an FDR close to 0.05. Therefore, we chose Lsize=5 for all analyses. To evaluate the robustness of this choice, we evaluate the similarity of the identified TAD structures between Lsize=5 and Lsize=6-10, and found that they are similar, with the median of the adjusted rand indices >0.75 (Supplementary figure 10). It indicates the result is relatively insensitive to the value of Lsize when Lsize=5∼10.

### Computation complexity of the TAD calling algorithm

We performed an analysis on the computational complexity for our recursive algorithm. For an *l*x*l* Hi-C matrix, if all bins are potential boundaries, then the recursion needs to visit *l*(*l*+1)/2 diagonal block sub-matrices. As there are *l* size 1 diagonal block matrices, the computation complexity for computing the scores of all size 1 matrices is O(*l*). Given the scores of size 1 matrices, we can calculate the scores of size 2 matrices. There are (*l*−1) of them, each enumerating through (2-1) partitions. Hence the time complexity is O((2-1)(*l*−1)). Following the same calculation, the scores of one sub-matrix of size *k* will be computed by enumerating (*k*−1) partitions. As there are (*l*-*k*+1) of them, the time complexity is O((*k*−1)(*l*-*k*+1)). Similar calculation can be done for the mean of sub-matrices. As a result, the total complexity to obtain the scores of all sub-matrices from size 1 to *l* is O(*l*^*3*^).

Empirically, the computational complexity is much lower than the above due to some further reductions. First, because potential TAD boundaries are limited to the TOPDOM local minimums, this substantially reduces the number of partitions from O(*l*^*3*^) to O(*m*^*3*^), where *m* is the number of candidate boundaries. Second, because TADs usually are smaller than 2Mb, the maximum TAD size to be called (*d*) typically is much smaller than *l*. This constraint effectively reduces the time complexity of our algorithm from O(*m*^*3*^) to O(*md*^*2*^). Furthermore, because TADs usually are formed between neighboring boundaries, we set a constraint in the recursive procedure to limit the TADs to be formed only between candidate boundaries that are no more than five neighbors apart.

### TAD-adjR^2^ for assessing accuracy of TAD calling

Because TADs are regions with frequent local interactions, a reasonable TAD caller is expected to classify the regions with high contact frequencies as TADs and the regions with low contact frequencies as non-TADs, i.e. gaps between TADs. At any given genomic distance, the variation between Hi-C signals should be largely explained by the classification of TADs. How well the variation can be explained by the classification of TADs can reflect the accuracy of TAD calling. Based on this intuition, we developed a metric similar to the R-square in regression models to evaluate the accuracy of TAD calling. Let *Y*_*i*_ denote the contact frequency of the *i*th bin, *n* denote the number of bins at the same genomic distance as this bin, *p* denotes the number of called TADs whose sizes are greater than or equal to the genomic distance. For bins within TAD, Ŷ_ℓ_denotes the average contact frequency at given genomic distance within that TAD, excluding regions covered by higher level TADs. For those bins not in any TADs, Ŷ_ℓ_ is the average of contact frequency in the gap region at that genomic distance. And 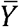 denote the overall mean contact frequency across all the bins at a given genomic distance. For each genomic distance, the TAD-adjR^2^ is defined as

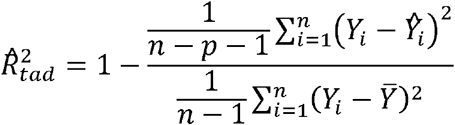

This quantity essentially measures the proportion of variance in Hi-C signal that is explained by the classification of TADs, adjusting for the number of TADs and genomic distance.

### Enrichment of expressed genes

To evaluate the activity of gene expression, we downloaded the RNA-seq data from ENCODE (See Data), merged the biological replicates of RNA-seq data, and computed the average FPKM for each gene. Genes with FPKM > 5 were deemed as expressed genes. For each TAD level, we compute the density of expressed gene as the number of expressed genes per 10Kb. For TADs with nested structures, genes covered by the inner level TADs are excluded in the calculation of gene density for outer TADs.

### Enrichment of CTCF or cohesin protein signals

To compute the enrichment of CTCF (or cohesin protein) signals at the identified boundaries and their surrounding regions, we computed the average CTCF (or cohesin proteins) signals from ChIP-seq data at the identified boundaries and the bins within their 10bins flanking regions. The processed signals in bigwig file was used in this process.

### Epigenetic state enrichment

We downloaded the IDEAS segmentation (see Data), which segments the genome into 36 epigenetic states based on 10 epigenomic marks [23]. We used it to evaluate the enrichment of epigenetic state in the identified (sub)TADs and boundaries. Let *n*_*i*_ denote the total number of 200bp windows that have IDEAS-assigned epigenetic states at a TAD boundary *i*, and *n*_*s,I*_ denote the number of 200bp windows annotated as state *s* at a TAD boundary *i*. For a given state *s,* its enrichment in a set of *M* boundaries is computed as

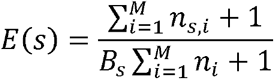

where *B*_*s*_ is the proportion of state *s* in the whole genome. The 1’s in the formula of E(s) are added to avoid dividing by 0.

### A/B compartments calling

We used CscoreTool [39] to infer A/B compartments from mouse G1E-ER4 Hi-C data (10Kb resolution, default parameter). The A/B compartments is determined by correlation coefficient between compartment score and ATAC-seq signal. If a positive correlation coefficient is observed, then regions with score>0 are in compartment A. Otherwise, if the correlation coefficient is below 0, the regions with score < 0 are in compartment A. We reversed the compartment scores on the chromosomes that has correlation coefficient < 0. Thus, compartment A is shown with positive score and compartment B is shown with negative score.

## Data

### Hi-C data

The human Hi-C data is obtained from Rao et al. 2014 (GEO accession number: GSE63525). Among them, three cell types (B-lymphoblastoid cells (GM12878), umbilical vein endothelial cells (HUVEC) and erythrocytic leukemia cells (K562)) were included in this study. The normalized (by Knight-Ruiz balancing method) Hi-C matrices at 5Kb, 10Kb and 25Kb resolutions were used in this study. The mouse Hi-C data is obtained from Hsu et al. 2017 [32](GEO accession number: GSE95476). The 10Kb raw Hi-C matrices from G1E-ER4 and two Brd2 knockouts were used in this study.

### Transcriptomic data

The gene expression data were downloaded from the ENCODE project (https://www.encodeproject.org/). The processed signal (FPKM) was used to measure the expression activity.

### Epigenomic data

The histone modification data were downloaded from the NIH Roadmap Epigenomics project (http://www.roadmapepigenomics.org/), including H2A.Z, H3K27ac, H3K27me3, H3K36me3, H3K4me1, H3K4me2, H3K4me3, H3K79me2, H3K9ac, H3K9me3 and H4K20me1. The ChIP-seq data of CTCF and cohesin protein (Rad21 and Smc3) were downloaded from ENCODE project (https://www.encodeproject.org/). The downloaded data were in BigWig format. The ‘bigWigAverageOverBed’ was used to segment signal into windows according to the resolution of Hi-C data. The mouse ChIP-seq data were downloaded from the VISION project (http://www.bx.psu.edu/~giardine/vision/).

### Epigenetic states

The IDEAS segmentation of the 6 ENCODE cell type/tissues (GM12878, H1h-ESC, Hela-S3, HepG2, HUVEC, K562) was downloaded from (http://main.genome-browser.bx.psu.edu/). The 36-state IDEAS model trained on 10 marks (H3K4me1, H3K4me2, H3K4me3, H3K9ac, H3K27ac, H3K27me3, H3K36me3, H3K20me1, PolII and CTCF), as well as DNase-seq and Faire-seq, was applied to this study.

---

**Supplementary figure 1.**
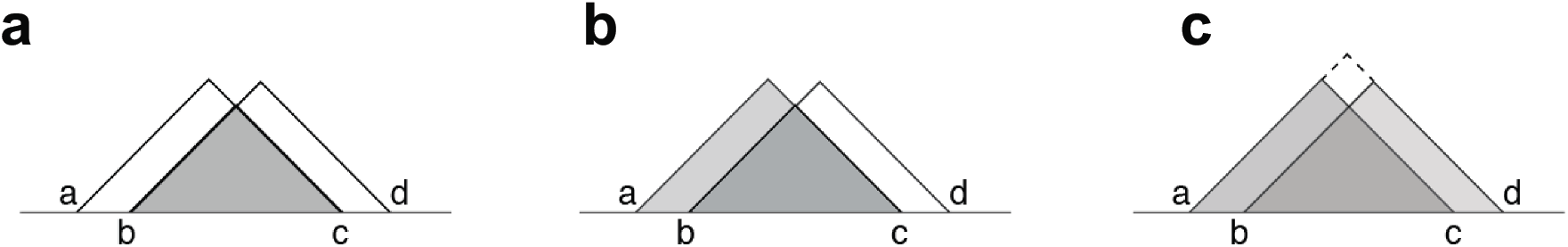
Illustration of convoluted TAD structures. **a**, Candidate TADs (a,c) and (b,d) are both suboptimal, as their scores may be driven by a real TAD (b,c). **b**, Two real TADs (a,c) and (b,c) are nested, which makes the score of (a,c) convoluted with the score of (b,c). **c**, Real TADs (a,c) and (b,d) are partially overlapping, which may be recaptured as nested TADs (b,c), (a,c) and (a,d).

**Supplementary figure 2.**
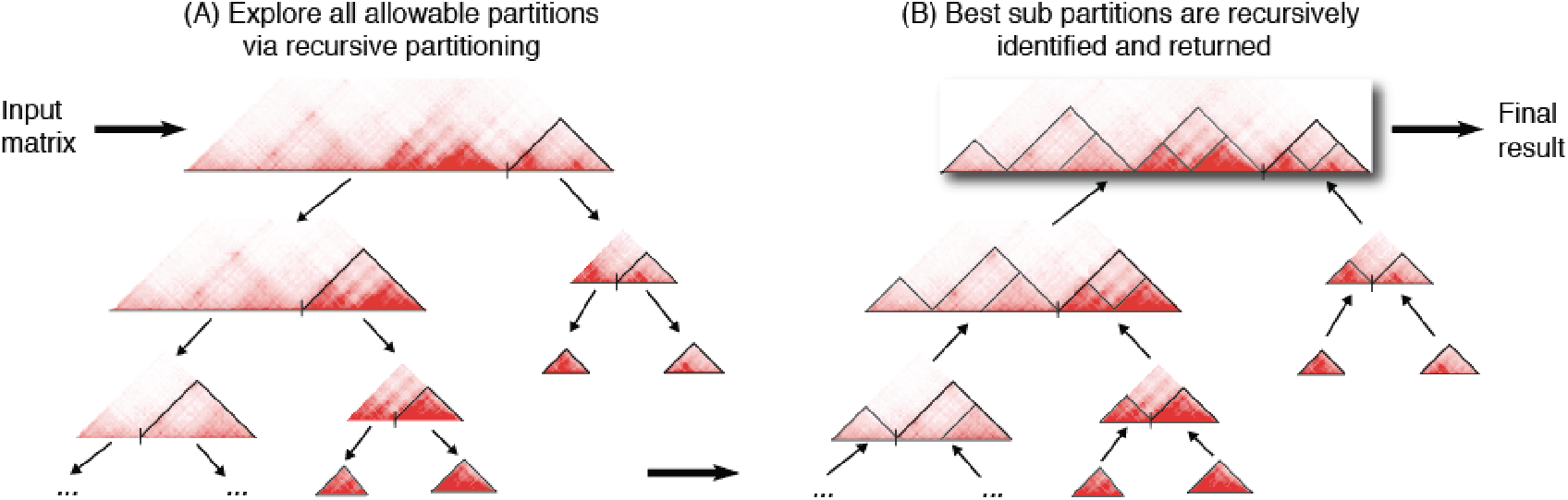
Illustration of the recursive TAD calling algorithm. **a**, At the first step of the algorithm, the entire Hi-C matrix is partitioned into two matrices, the one forming the largest right-most TAD (i.e. triangles marked in black) and the remaining part, according to a score function. Then the same function is called on each sub-matrix to recursively identify nested TAD structures. **b**, Each recursion step identifies the best set of TADs in its matrix under consideration according to the score function, and returns the TAD calls back to its parent until the root.

**Supplementary figure 3.**
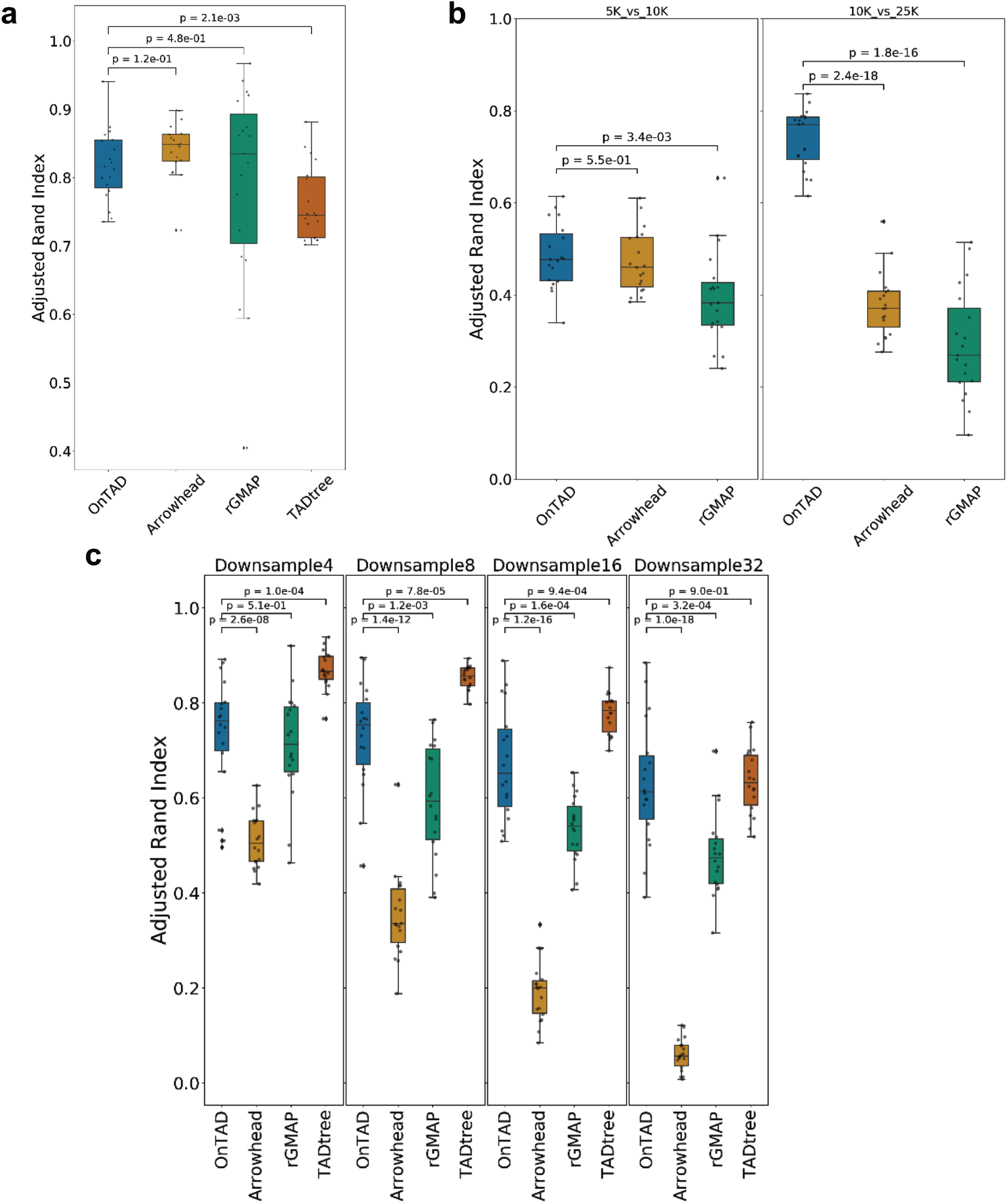
TAD reproducibility under different measurements. **a**, Adjusted rand index between TADs from two biological replicates (GM12878, 10Kb). **b**, Adjusted rand index across TADs from Hi-C data in multiple resolutions (GM12878, 5Kb, 10Kb and 25Kb). TADtree is not included because it has difficulty finishing the computation on high resolution data due to its high memory consumption. **c**, Adjusted rand index between TADs from Hi-C data in original sequencing depth and in different down sampled sequencing depth (GM12878, 1/4, 1/8, 1/16 and 1/32 of original sequencing depth).

**Supplementary figure 4.**
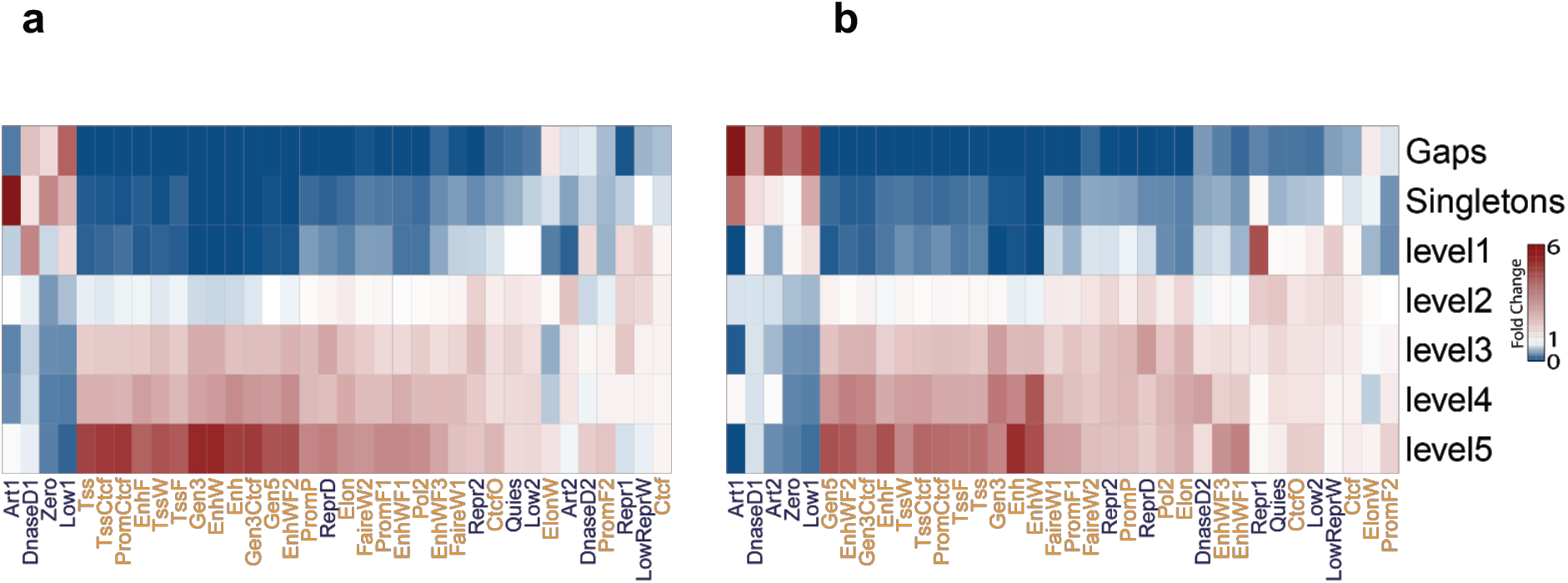
Enrichment of epigenetic states at the boundaries of different levels of TADs. Enrichment of epigenetic states at the regions covered by different levels of TADs. The enrichment (fold change) of active states (orange states) increases as the TAD level increases. **a**, K562 **b**, Huvec

**Supplementary figure 5.**
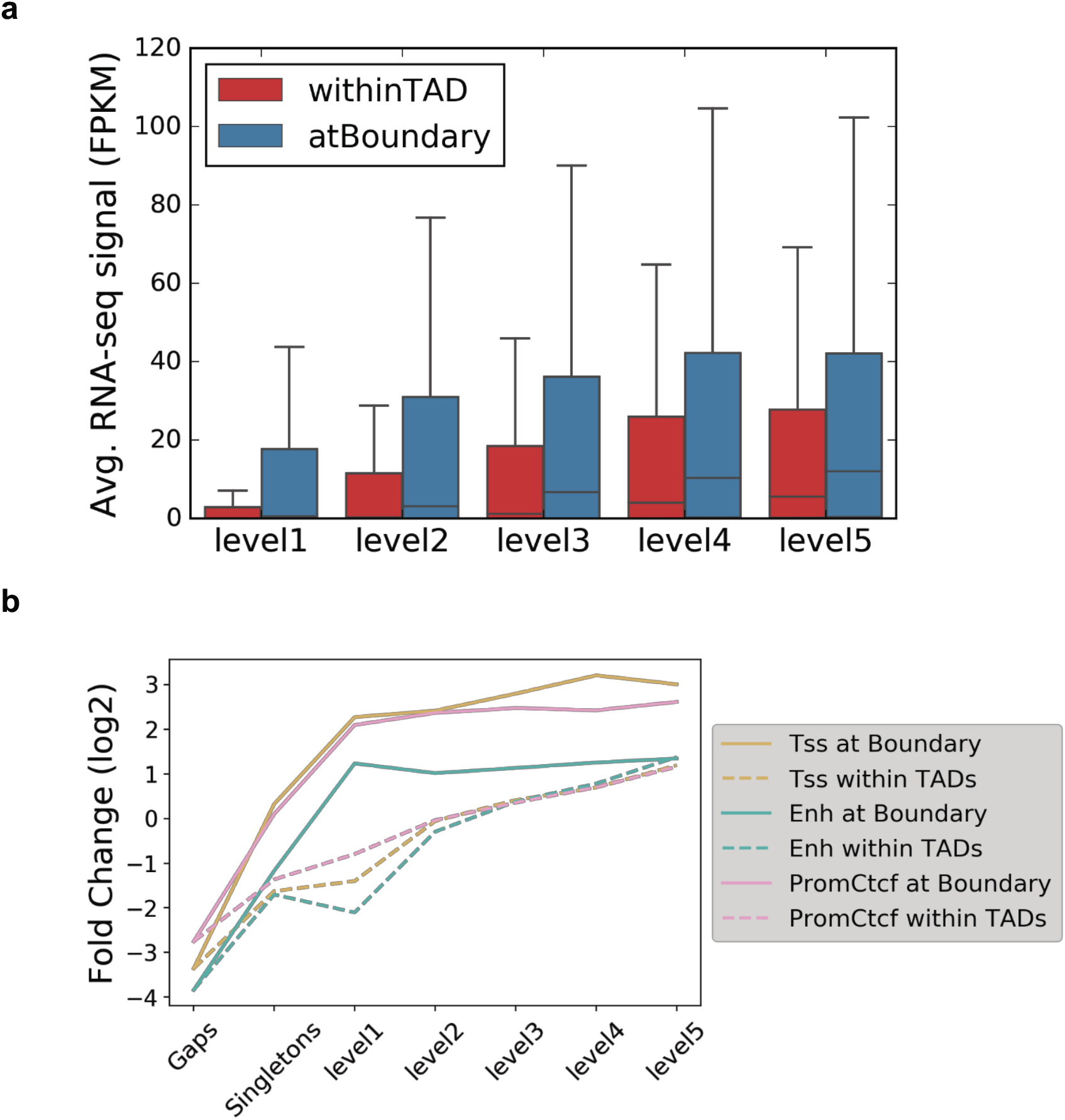
Comparison between boundaries and inside TADs. **a**, Distribution of RNA-seq signal (FPKM) at the boundaries (blue) and within TADs (red) **b**, Enrichment of active epigenetic states at the TAD boundaries (solid line) versus inside TADs (dashed line). Y-axis denotes fold enrichment of three active epigenetic states (Tss, Enh and PromCtcf). X-axis denotes the boundaries and TADs at different levels.

**Supplementary figure 6.**
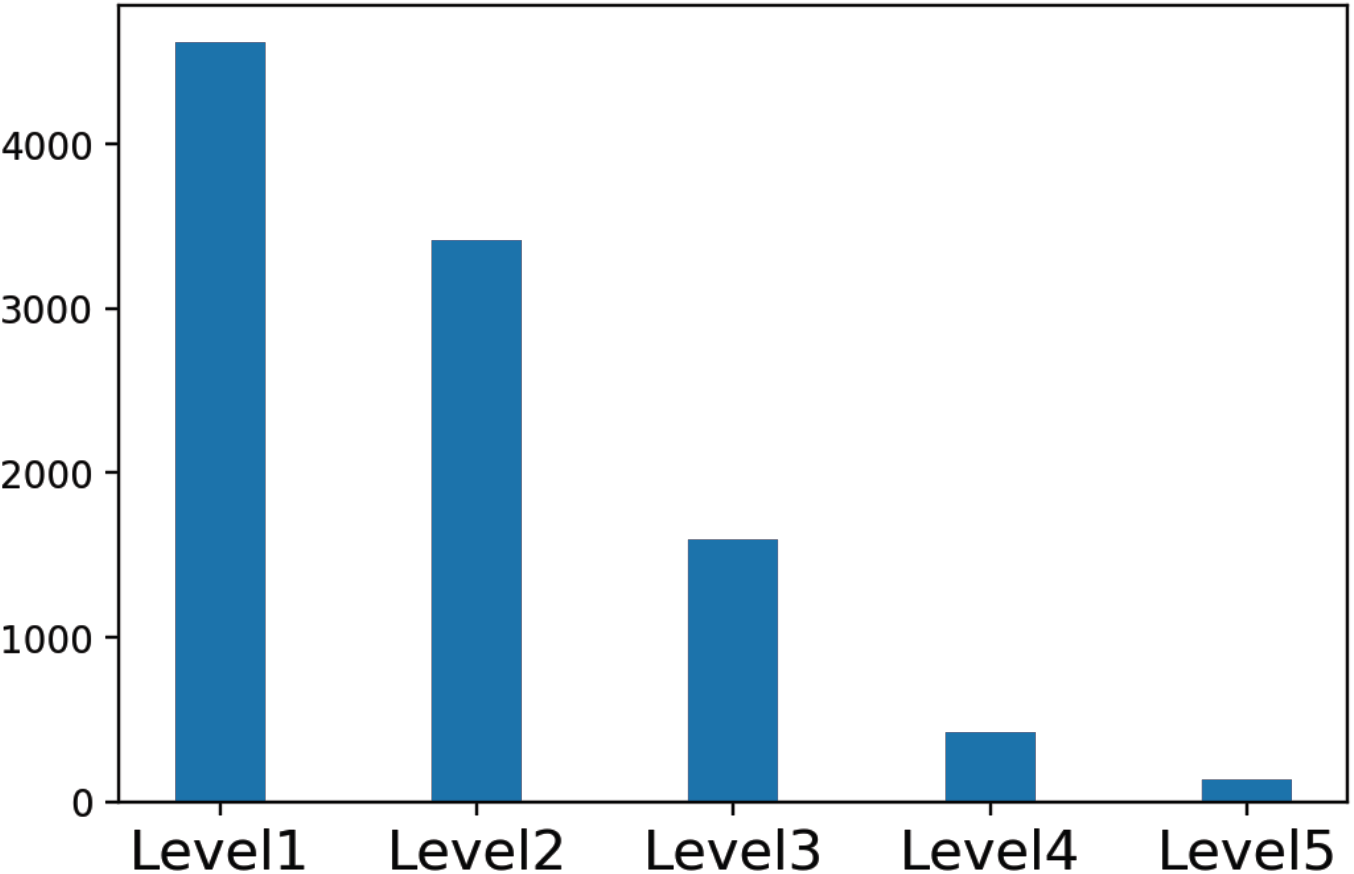
Distribution of the levels of TAD boundaries.

**Supplementary figure 7.**
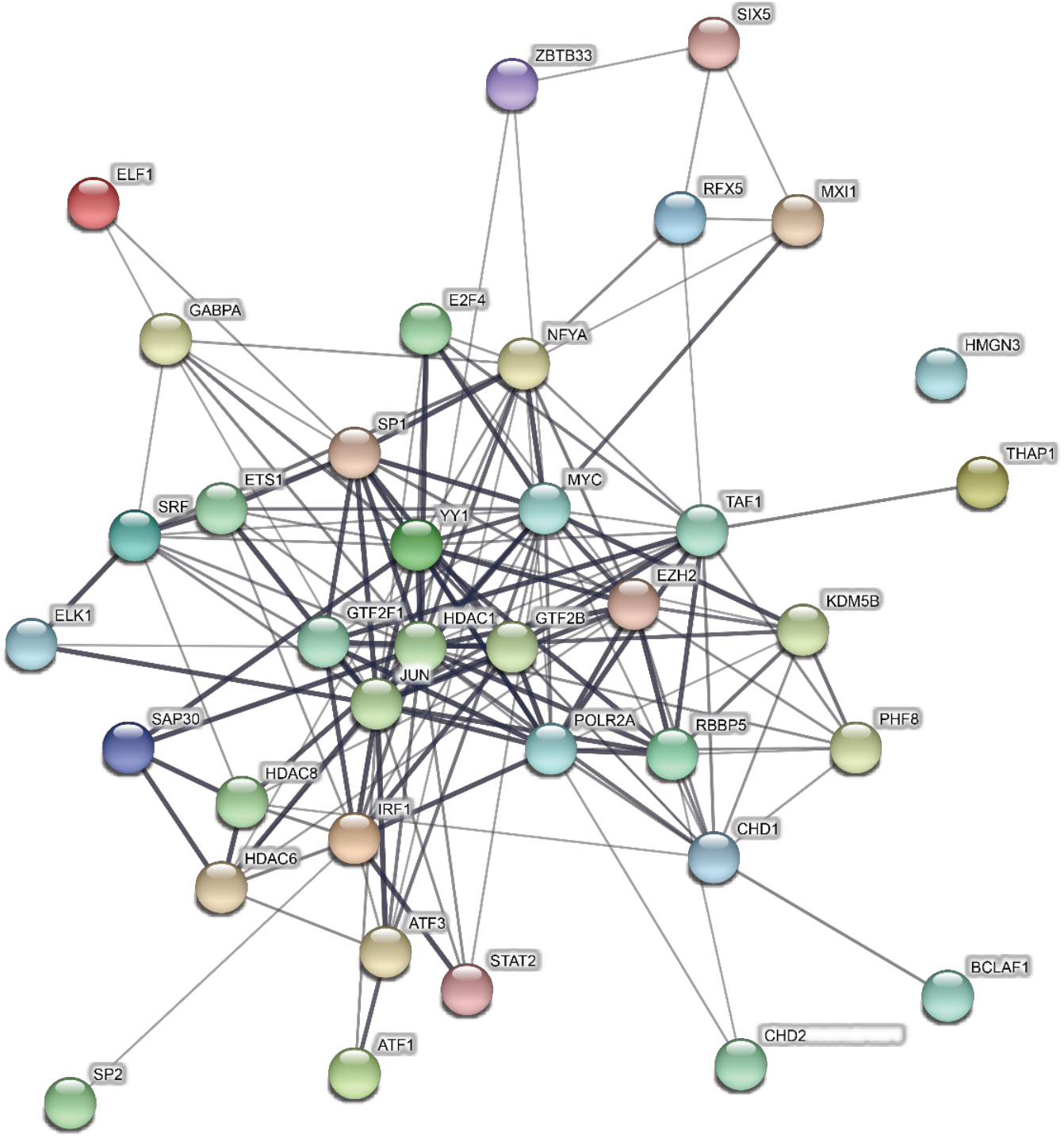
Protein-protein interaction network of hub-boundary-enriched TFs from STRING database. Each node denotes a TF that are at least 2-fold enriched at hub-boundary over level1 boundary (n=37). Each edge denotes the interaction potential between two TFs, with thicker edges corresponding to higher interaction confidence. Interaction data was downloaded from STRING database (https://string-db.org/)

**Supplementary figure 8.**
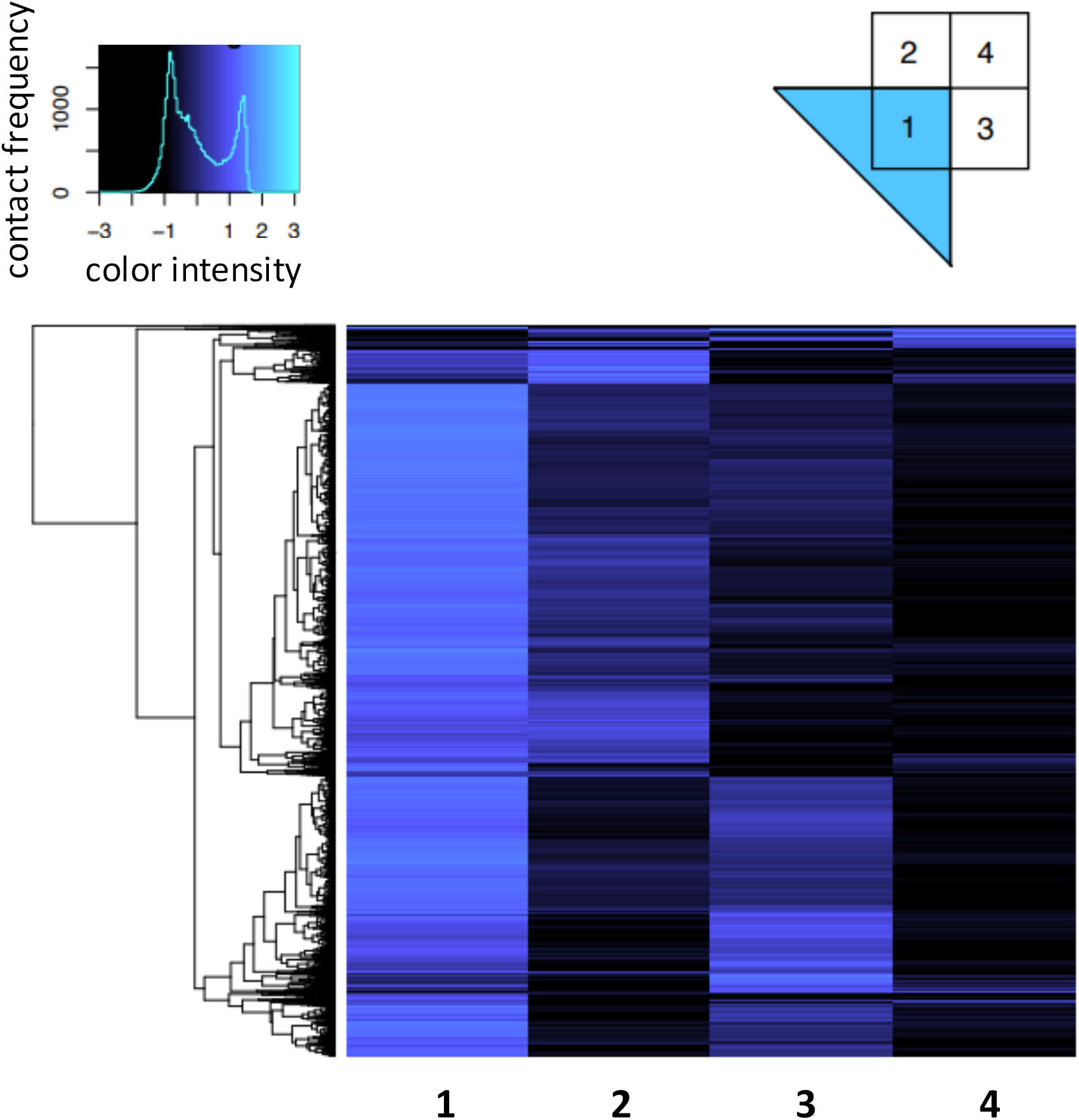
Contact frequency is unbalanced between the two sides of hierarchical TAD corners. The regions around TAD corners are segregated into four quadrants (1-4 on the top right figure). We then averaged contact frequency of each TAD corner by quadrants. As shown in the heatmap, the majority quadrant 2 and 3 shows unequal average contact frequencies, suggesting that the inner TADs tend to be formed on one side of the outer TADs, rather than on both sides. Quadrant 1 has the highest average contact frequency because it is within TADs.

**Supplementary figure 9.**
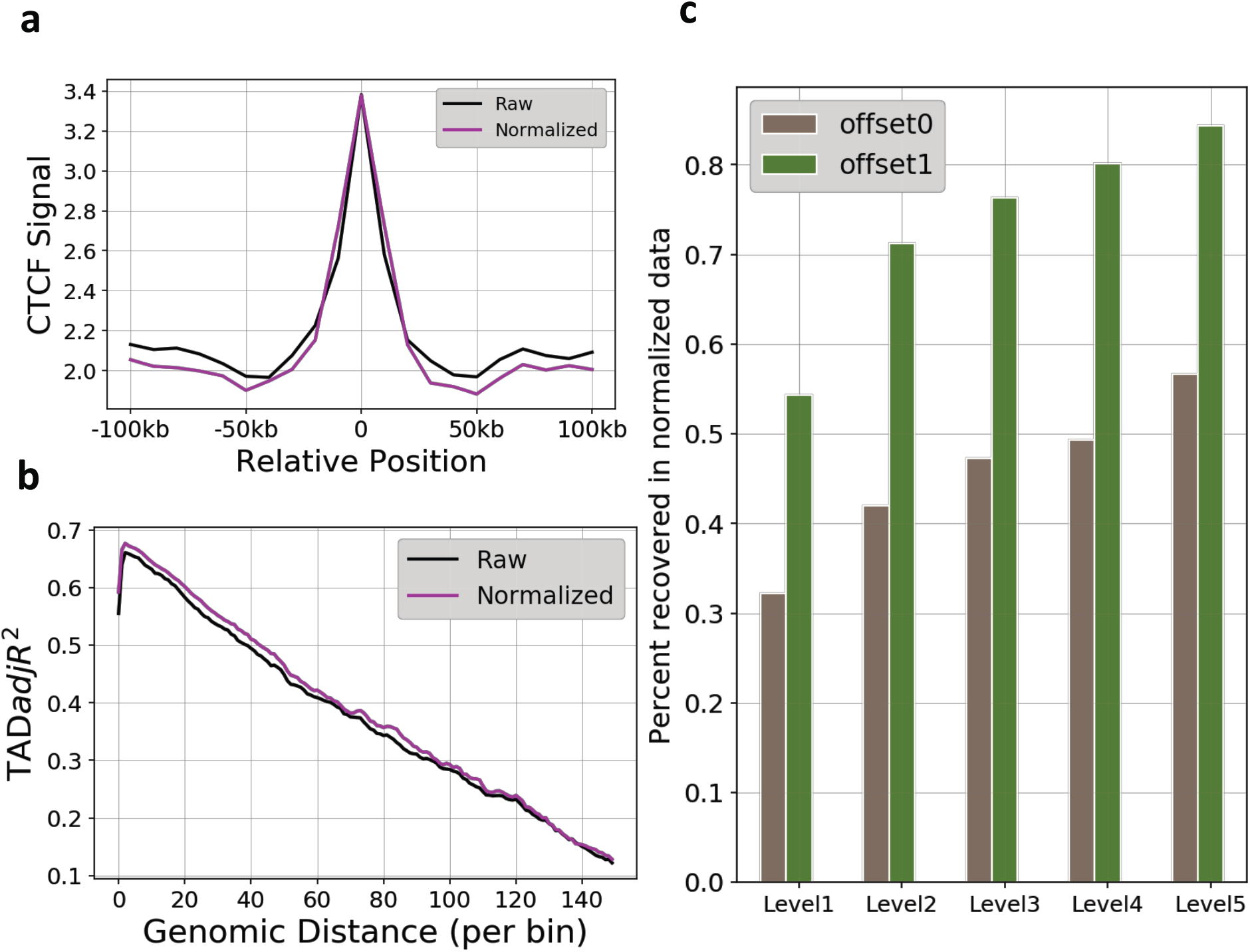
Comparison of OnTAD results between raw Hi-C and normalized Hi-C in GM12878 (10kb). **a**, Enrichment of CTCF signal at identified TAD boundaries and surrounding regions (+/− 10 bins) in raw Hi-C matrix and normalized Hi-C matrix. Y-axis: The average ChIP-Seq signal. **b**, TAD-adjR^2^ of OnTAD results at difference genomic distance in raw Hi-C matrix and normalized Hi-C matrix. The results on normalized data show a slightly higher enrichment of CTCF at boundary and a higher TAD-adjR^2^. The normalized Hi-C matrices were generated by Knight-Ruiz balancing [40] method. **c**, The proportion of boundaries identified in raw data that are recovered in normalized data. Grey: exact match; Green: one bin offset allowed when matching the boundaries identified in raw and normalized data. Half of the boundaries identified in the raw data precisely match with the boundaries identified in the normalized data. If we allow one bin offset when matching the locations of the boundaries, over 71% of the high-level TAD boundaries (level 2+) are matched between the results from raw data and normalized data.

**Supplementary Figure 10.**
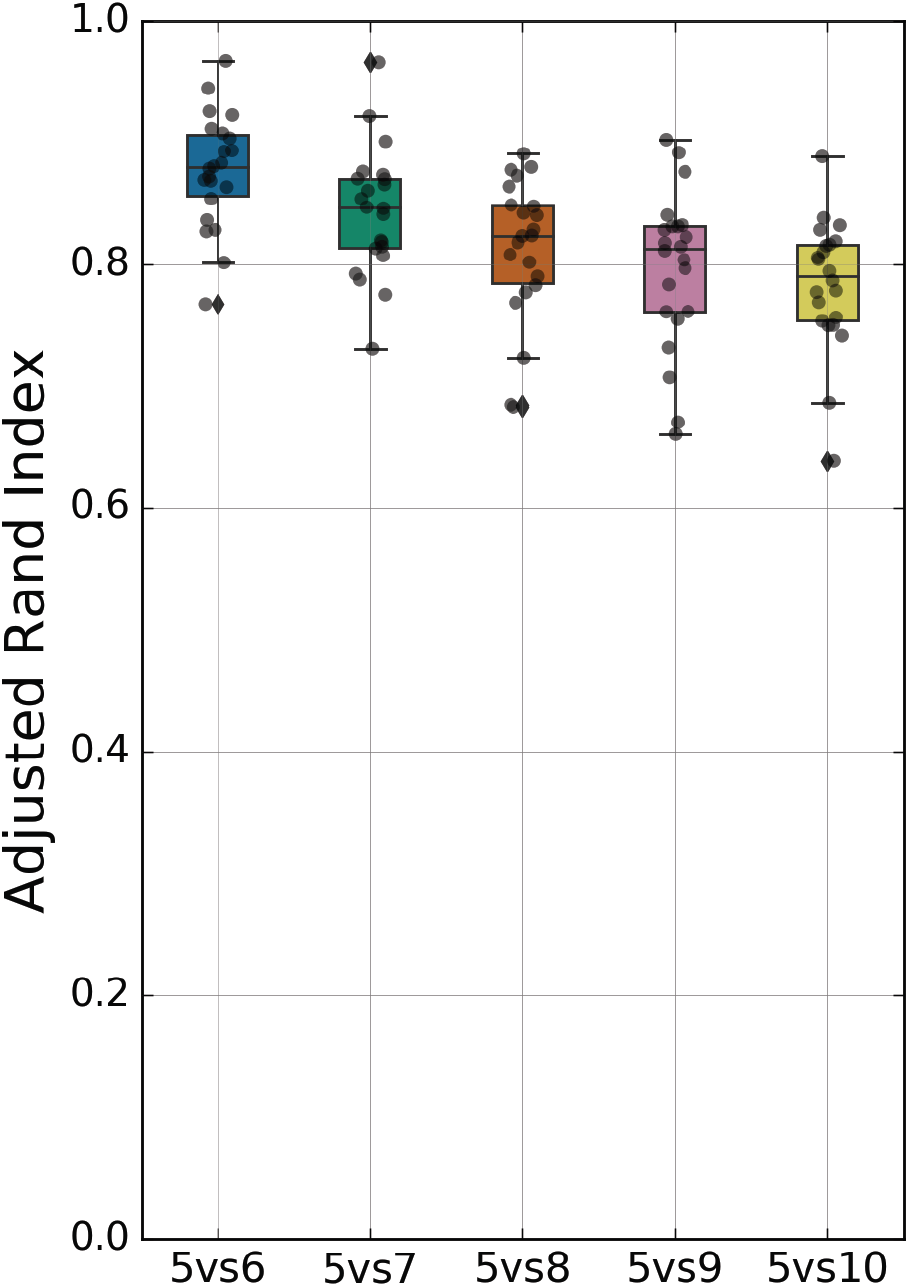
Similarity between the (sub)TADs identified at Lsize = 5 and at other Lsizes.

**Supplementary Table 1 | Comparison of running time of different methods on high resolution Hi-C data (GM12878 10Kb) (unit: seconds)**.

**Supplementary Table 2 | Number of TADs on each side of a boundary that share this boundary (GM12878 10Kb)**.

**Supplementary Table 3 | The FDR and number of TADs under each penalty value. (GM12878, average on 100 permutations)**

**Supplementary Table 4 | The FDR and number of TADs under each penalty value. (G1E-ER4, average on 100 permutations)**

**Supplementary Table 5 | The FDR and number of TADs under each Lsize. (GM12878, average on 100 permutations)**

**Supplementary File1 | Commands for operating other TAD calling methods.**

## Declarations

### Acknowledgments

This work was supported by the NIH R01 GM121613, NIH R01GM109453 and NIH R24 DK106766, NIH training grant T32 GM102057 (CBIOS training program to The Pennsylvania State University), a Huck Graduate Research Innovation Grant, and 4D Nucleome NIH Initiative DK107980 (Center for 3D Structure and Physics of the Genome).

### Availability of data and materials

OnTAD is available at https://github.com/anlin00007/OnTAD.

### Authors’ contributions

YZ and LA implemented OnTAD. YZ and QL conceived the method. LA, TY, GX and JY conducted the analysis. LA, YZ, QL, TY and RCH wrote the manuscript with assistance from the other authors. JN assisted the interpretation of the results. All authors read and approved the final manuscript.

