## Supplementary Tables for "Hierarchical Domain Structure Reveals the Divergence of Activity among TADs and Boundaries"

**Supplementary table 1: Comparison of running time of different methods on high resolution Hi-C data (GM12878 10Kb) (unit: seconds).**

|  | OnTAD | Arrowhead | rGMAP | Dicaller | TADtree* |
| --- | --- | --- | --- | --- | --- |
| chr1 | 83 | 225 | 2902 | 1669 |  |
| chr2 | 73 | 234 | 1563 | 1463 |  |
| chr3 | 59 | 174 | 1215 | 1018 |  |
| chr4 | 55 | 168 | 1139 | 1023 | 56742 |
| chr5 | 46 | 161 | 1015 | 1111 | 44751 |
| chr6 | 41 | 153 | 1069 | 1057 | 26086 |
| chr7 | 36 | 105 | 832 | 925 | 14708 |
| chr8 | 32 | 98 | 789 | 739 | 9270 |
| chr9 | 30 | 91 | 2085 | 658 | 8997 |
| chr10 | 27 | 94 | 721 | 851 | 8365 |
| chr11 | 28 | 91 | 704 | 839 | 8368 |
| chr12 | 27 | 87 | 729 | 765 | 8166 |
| chr13 | 22 | 82 | 443 | 578 | 6874 |
| chr14 | 17 | 74 | 403 | 469 | 6584 |
| chr15 | 16 | 72 | 782 | 328 | 5984 |
| chr16 | 13 | 63 | 387 | 463 | 5363 |
| chr17 | 12 | 67 | 323 | 444 | 4119 |
| chr18 | 11 | 57 | 337 | 429 | 3926 |
| chr19 | 8 | 38 | 222 | 246 | 2896 |
| chr20 | 8 | 40 | 487 | 304 | 3065 |
| chr21 | 5 | 30 | 153 | 279 | 2417 |
| chr22 | 6 | 32 | 190 | 183 | 2559 |
| Total | 655 | 2236 | 18490 | 15841 | 172498 |

***Due to the limit of walltime (up to 120hrs) and computing resources (up to 120Gb), we cannot finish running of TADtree on chr1, chr2 and chr3**

**Supplementary table 2: Number of TADs on each side of a boundary that share this boundary (GM12878 10Kb).**

| Num of TADs on left\right | 0 | 1 | 2 | 3 | 4 | 5 |
| --- | --- | --- | --- | --- | --- | --- |
| 0 | 0 | 1297 | 378 | 110 | 23 | 9 |
| 1 | 1209 | 2108 | 1060 | 375 | 77 | 19 |
| 2 | 428 | 1023 | 524 | 214 | 80 | 27 |
| 3 | 100 | 413 | 247 | 136 | 41 | 7 |
| 4 | 22 | 71 | 61 | 35 | 12 | 3 |
| 5 | 7 | 24 | 17 | 17 | 5 | 1 |

**Supplementary table 3:** **The FDR and number of TADs under each penalty value. (GM12878, average on 100 permutations)**

| Penalty (λ) | FDR | # of TADs |
| --- | --- | --- |
| **0.0** | 0.089 | 15058 |
| **0.1** | 0.054 | 13236 |
| **0.2** | 0.041 | 11371 |
| **0.3** | 0.020 | 9461 |
| **0.4** | 0.015 | 7848 |
| **0.5** | 0.011 | 6394 |

**Supplementary table 4: The FDR and number of TADs under each penalty value. (G1E-ER4, average on 100 permutations)**

| Penalty (λ) | FDR | # of TADs |
| --- | --- | --- |
| **0.0** | 0.083 | 9025 |
| **0.1** | 0.028 | 7586 |
| **0.2** | 0.017 | 6303 |
| **0.3** | 0.013 | 5155 |
| **0.4** | 0.005 | 4230 |
| **0.5** | 0.005 | 3383 |

**Supplementary table 5: The FDR and number of TADs under each Lsize. (GM12878, average on 100 permutations)**

| Lsize | FDR | # of TADs |
| --- | --- | --- |
| **3** | 0.058 | 12987 |
| **4** | 0.070 | 13579 |
| **5** | 0.055 | 13236 |
| **6** | 0.053 | 12626 |
| **7** | 0.037 | 11920 |
| **8** | 0.036 | 11287 |
| **9** | 0.038 | 10677 |
| **10** | 0.038 | 10147 |
